# Reliable and efficient parameter estimation using approximate continuum limit descriptions of stochastic models

**DOI:** 10.1101/2022.02.02.478913

**Authors:** Matthew J. Simpson, Ruth E. Baker, Pascal R. Buenzli, Ruanui Nicholson, Oliver J. Maclaren

## Abstract

Stochastic individual-based mathematical models are attractive for modelling biological phenomena because they naturally capture the stochasticity and variability that is often evident in biological data. Such models also allow us to track the motion of individuals within the population of interest. Unfortunately, capturing this microscopic detail means that simulation and parameter inference can become computationally expensive. One approach for overcoming this computational limitation is to coarse-grain the stochastic model to provide an approximate continuum model that can be solved using far less computational effort. However, coarse-grained continuum models can be biased or inaccurate, particularly for certain parameter regimes. In this work, we combine stochastic and continuum mathematical models in the context of lattice-based models of two-dimensional cell biology experiments by demonstrating how to simulate two commonly used experiments: cell *proliferation assays* and *barrier assays*. Our approach involves building a simple statistical model of the discrepancy between the expensive stochastic model and the associated computationally inexpensive coarse-grained continuum model. We form this statistical model based on a limited number of expensive stochastic model evaluations at design points sampled from a user-chosen distribution, corresponding to a computer experiment design problem. With straightforward design point selection schemes, we show that using the statistical model of the discrepancy in tandem with the computationally inexpensive continuum model allows us to carry out prediction and inference while correcting for biases and inaccuracies due to the continuum approximation. We demonstrate this approach by simulating a *proliferation assay*, where the continuum limit model is the well-known logistic ordinary differential equation, as well as a *barrier assay* where the continuum limit model is closely related to the well-known Fisher-KPP partial differential equation. We construct an approximate likelihood function for parameter inference, both with and without discrepancy correction terms. Using maximum likelihood estimation, we provide point estimates of the unknown parameters, and use the profile likelihood to characterise the uncertainty in these estimates and form approximate confidence intervals. For the range of inference problems considered, working with the continuum limit model alone leads to biased parameter estimation and confidence intervals with poor coverage. In contrast, incorporating correction terms arising from the statistical model of the model discrepancy allows us to recover the parameters accurately with minimal computational overhead. The main tradeoff is that the associated confidence intervals are typically broader, reflecting the additional uncertainty introduced by the approximation process. All algorithms required to replicate the results in this work are written in the open source Julia language and are available at GitHub.

## 1 Introduction

Discrete, stochastic models of cell biology phenomena are often used to simulate a range of twodimensional cell biology experiments, including *cell proliferation assays* [4, 5, 50] and *cell migration assays* [29, 49]. A cell proliferation assay is extremely simple, and involves uniformly seeding a population of cells, at low density, on a two-dimensional cell culture plate [4, 5, 50]. Over a short time scale of a few hours, cells attach to the plate and migrate, often randomly. Over a longer time scale of 24-48 hours, cells proliferate, and the result of the combined cell migration and cell proliferation is the formation of a dense monolayer, as depicted in the simulation of a proliferation assay in Figure 1(a). By reporting the increase in monolayer density over time, data from cell proliferation assays can be used to estimate the proliferation rate and doubling time of the cell population [5,50,54]. A cell migration assay is slightly more complicated since cells are initially distributed non-uniformly on a two–dimensional cell culture plate, and there are two main ways of achieving this. One approach, called a scratch assay, involves seeding a population of cells on a two-dimensional cell culture plate, allowing a certain duration of time for the monolayer density to increase, and then scratching the monolayer using a fine-tipped instrument [29, 56]. Another approach, which avoids scratching the monolayer, is called a *barrier assay* where cells are placed onto the two-dimensional cell culture plate inside a physical barrier [8, 16]. This procedure creates an initially-confined population of cells that tends to spread outwards after the barrier is lifted. Both barrier and scratch assays involve some area of the cell culture plate being occupied by a monolayer of cells whereas adjacent areas are completely vacant. Over time, combined cell migration and proliferation leads to the initially-confined population spreading into the previously-vacant parts of the cell culture plate, as depicted in the simulation of a barrier assay in Figure 1(b) [8,16]. Observations of collective cell spreading and cell invasion process are used to quantify rates of cell migration and cell proliferation [16, 35, 36, 56].

**Figure 1:**
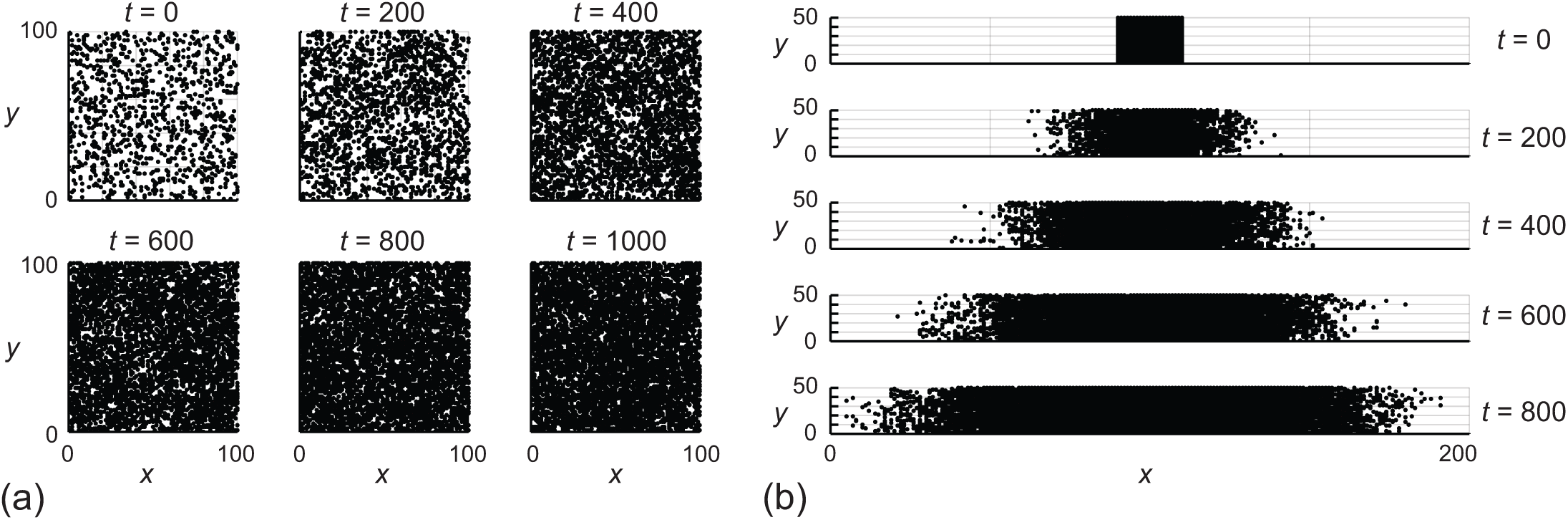
(a) Discrete snapshots of the evolution of the stochastic model to simulate a cell proliferation assay on a lattice with *I* = *J* = 100 and *C*(0) = 0.1, *R*_*m*_ = 1, *R*_*p*_ = 0.01 and *R*_*d*_ = 0.005 at *t* = 0, 200, 400, 600, 800 and 1000. (b) Discrete snapshots of the evolution of the stochastic model to simulate a barrier assay on a lattice with *I* = 200 and *J* = 50, with *C*_*i*_(0) = 1 for *i* ∈ [91, 111] and *C*_*i*_(0) = 0 elsewhere. The evolution of the simulation corresponds to *R*_*m*_ = 1, *R*_*p*_ = 0.01 and *R*_*d*_ = 0 at *t* = 0, 200, 400, 600 and 800.

There are several advantages of modelling these kinds of experiments with a discrete, stochastic modelling framework [38, 61]. First, a discrete stochastic modelling approach allows us to model individual-level behaviour and captures variability [5,31,32,34]. Second, a discrete stochastic modelling approach can produce snapshots of the experiments that are consistent with the kinds of images that are reported experimentally [5, 31, 32, 34]. The key limitation of working with a stochastic modelling approach relates to computational requirements since the computation time increases with the number of individuals simulated and the complexity of how the individuals interact. Using a discrete stochastic modelling framework for parameter inference can be very challenging, especially in a Bayesian framework that might require millions of expensive simulations to be performed [53, 58]. The computational bottleneck associated with discrete stochastic models is even more pressing if we are interested in spatiotemporal processes, such as modelling barrier assays and scratch assays, rather than simpler models of well-mixed stochastic processes, since here we wish to keep track of agent position and this incurs additional computational cost that is avoided in simulations of well-mixed phenomena.

A different approach to modelling cell biology experiments is to use continuum mathematical models. One way to obtain such a model is by coarse-graining the kinds of discrete stochastic models discussed previously and illustrated in Figure 1 [49]. A key advantage of working with a coarse-grained differential equation model is that the computational effort required to solve the governing equation is typically less than the computational effort required to simulate a stochastic model. Furthermore, the computational effort required to solve a coarse-grained differential equation model is independent of the population size. Together, these two features mean that parameter inference is computationally tractable. One limitation of working with a coarse-grained differential equation model, however, is that these models do not predict individual-level behaviour, nor do they capture variability in the way that stochastic models do. A more important limitation, however, is the fact that coarse-grained differential equation models are generally only accurate in fairly restricted regions of the parameter space [1, 51]. For example, in the simulation of barrier assays and proliferation assays, continuum-limit descriptions of discrete exclusion process-based stochastic models are accurate only in the limit where the rate of proliferation is very small compared to the rate of migration [49]. While some attention has been given to developing alternative continuum-limit descriptions that include describing pair-wise interactions [2, 3, 6, 13, 27], these alternative continuum-limit descriptions often involve relatively complicated moment closure approximations, and it is often unclear which kind of closure approximation should be used since different approximations can lead to different outcomes. In addition, introducing a pair-wise framework and moment closure approximations can improve accuracy [1], this improvement comes at the expense of incurring substantial increases in computational overhead [1]. Furthermore, and most importantly, just like standard coarse-grained differential equation descriptions, computationally expensive moment closure-based approximations also fail to accurately predict data from stochastic simulation algorithms in certain parameter regimes [51]. While the focus of this work is to examine lattice-based models of cell biology experiments, it is well-known that coarse-grained descriptions of other types of stochastic models of biological phenomena can also fail to capture the underlying stochastic process for certain parameter choices [30, 55].

Here we demonstrate a different approach to deal with prediction and inference that combines the computational simplicity of a continuum differential equation model with the accuracy of a stochastic modelling framework for modelling cell biology experiments. This approach takes advantage of having a model family that includes both an accurate (fine) discrete stochastic simulation model and an associated approximate (coarse) continuum limit model [49]. In particular, we show how to use the coarse continuum limit model for the primary, computationally expensive inference and prediction steps while incorporating an additional statistical ‘meta-model’ [12, 15, 40] of the discrepancy between the two primary models. This meta-model allows us to correct for the bias and uncertainty that arises from the use of an approximate mechanistic process model. We build the discrepancy model before carrying out the inference and prediction steps using a finite number of paired fine and coarse model runs at ‘design points’ chosen from a user-specified distribution. This initial step enables careful control over the computational budget. Our straightforward construction of the discrepancy meta-model amounts to carrying out multivariate multiple linear regression of the (potentially spatiotemporal) discrepancy between the fine and coarse mechanistic models in terms of their shared parameters. Naturally, more complex meta-models are possible, but this approach is straightforward, and we show that it works well in practice. Furthermore, the availability of a coarse model with parameters in common with the fine model enables a relatively simple statistical model to be plausible – we only need to statistically model the discrepancy between the approximate- and fine-scale models rather than the fine model itself. In essence, our methodology corresponds to the ‘Bayesian approximation error’ (BAE) [20–22] approach to model errors, which we have used in other applications [33], though here we present it in a form compatible with both Bayesian and frequentist statistical inference.

More broadly, our work falls under the umbrella of ‘design of computer experiments’ and the associated topic of meta-modelling of expensive simulation models; such models are also known as ‘surrogates’ or ‘emulators’ [12, 15, 40]. There is extensive literature on this topic, with one of the first articles on computer experiments dating back to 1979 [37]. Early literature in this area focused mainly on the approximation of expensive deterministic models by statistical models, which leads to subtleties over interpretations of the randomness in the meta-models [25, 41]. Arguably, this issue led to much of the literature taking a Bayesian approach in which probability can naturally reflect ignorance rather than actual natural variation. Two popular methodologies in this style are the Kennedy and O’Hagan (KOH) [23, 24] and the BAE [20–22] approaches. The former uses infinite-dimensional Gaussian processes with a restricted form of the covariance function (potentially requiring hyperparameter estimation); the latter uses a finite-dimensional multivariate Gaussian approximation but an empirical (unconstrained) covariance matrix. As mentioned, our approach here is essentially the same as this latter (BAE) work, though with minor differences that we feel provide greater flexibility. In particular, our approach is also consistent with a frequentist viewpoint, like that developed by Owen [41] and Koehler and Owen [25] in the context of deterministic computer experiments where the randomness is strictly due to sampling variability of the design points. We emphasise the ability to choose flexible sampling plans for the design points, thus representing the design of experiments point of view rather than (necessarily) representing a prior distribution of belief. Furthermore, as we deal with a stochastic fine-scale model, many interpretational difficulties associated with statistically approximating a deterministic model disappear naturally.

Our approach of using a surrogate model to facilitate prediction and inference for a more accurate, but computationally expensive model falls within a wide class of approaches used across various disciplines such as mathematical models of: water resources and fluid flow in porous media [60]; biomechanics and bone deformation [62]; electrophysiology [28] and gene expression [18]. In some of these applications the coarse and fine models are simply taken to be different numerical discretizations of the same governing differential equations, which often involve multi-dimensional PDE models [60, 62]. Another approach to define coarse and fine models is to work with different mathematical models describing the same phenomena, but with different levels of complexity inherent in the modelling assumptions [18, 28]. Our approach is different in the sense that we use a standard coarse-graining approximation to relate a computationally expensive discrete stochastic mathematical model to a relatively inexpensive continuum limit differential equation-based approximation.

To illustrate our methodology, we describe and implement a standard on-lattice exclusion-based model of cell migration, cell proliferation and cell death and show how this discrete model can be used to mimic proliferation and barrier assays. We then derive the classical continuum-limit differential equation descriptions of the discrete models and show how the continuum limit descriptions accurately capture data from the discrete models for certain parameter choices whereas the same continuum limit descriptions lead to inaccurate predictions for other parameter choices. Next, we lay out a simple algorithm to build a discrepancy term that proves to be useful in making predictions (i.e. the forwards problem) and parameter inference (i.e. the backwards or inverse problem). After first demonstrating these ideas in the simplest case of a proliferation assay, we repeat all calculations in the more complicated setting of a barrier assay. All algorithms required to replicate the results in this work are written in the open-source Julia language and are available at GitHub.

## 2 Results and discussion

### 2.1 Discrete model and continuum limit derivation

Simulations are performed on a two-dimensional square lattice with lattice spacing Δ. Site are indexed in the usual way, *i, j* ∈ ℤ, so that each site has position (*x, y*) = (*i*Δ, *j*Δ). All simulations are performed on a lattice of size *I* ×*J*, where *i* = 0, 1, 2, …, *I* and *j* = 0, 1, 2, …, *J*, so that the total number of lattice sites is (*I* + 1)(*J* + 1). The lattice spacing can be thought of as being the size of a cell diameter [9, 53], and all analysis carries across directly to other regular lattices, such as a hexagonal lattice. In any single realisation of the discrete model the occupancy of site (*i, j*) is denoted *c*_*i,j*_, with *c*_*i,j*_ = 1 for an occupied site and *c*_*i,j*_ = 0 for a vacant site.

We evolve the discrete model through time using the Gillespie algorithm [14]. Agents migrate at rate *R*_*m*_, proliferate at rate *R*_*p*_ and die with rate *R*_*d*_. To assess a potential motility event for an agent at (*x, y*), a target site is chosen at random from one of the four nearest neighbour lattice sites at locations (*x* ± Δ, *y* ± Δ). Potential movement events are successful if the target site is vacant, otherwise the attempted movement event is aborted. To assess a potential proliferation event for an agent at (*x, y*), a target site for the placement of a daughter agent is chosen at random from one of the four nearest neighbour lattice sites at locations (*x* ± Δ, *y* ± Δ). If the target site is vacant a new agent is placed on the lattice. Agent death, in contrast, does not involve any movement or placement of daughter agents on the lattice. This means that agent death does not involve any assessment of the occupancy of nearest neighbour lattice sites. We implement the discrete model with reflecting boundary conditions along all boundaries, and we always work with dimensionless simulations by setting Δ = 1. The results can be re-scaled to match any particular experimental measurements by appropriate re-scaling of Δ [49].

To connect the discrete mechanism to a continuum model we average the occupancy of site (*i, j*) over many identically prepared realizations of the model to obtain ⟨*c*_*i,j*_⟩ ∈ [0, 1] [49]. After averaging we form a discrete conservation statement describing the time rate of change of ⟨*c*_*i,j*_⟩,

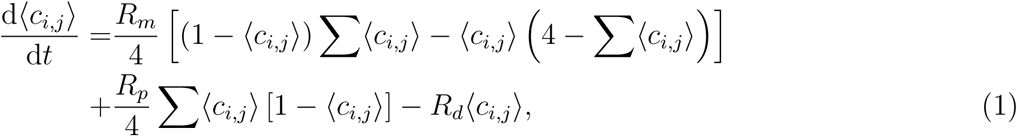

where Σ⟨*c*_*i,j*_⟩ = ⟨*c*_*i*+1,*j*_⟩ + ⟨*c*_*i*−1,*j*_⟩ + ⟨*c*_*i,j*+1_⟩ + ⟨*c*_*i,j*−1_⟩. The terms on the right of Equation (1) that are proportional to *R*_*m*_ describe the rate of change of average occupancy of site (*i, j*) due to net migration, whereas the remaining terms on the right of Equation (1), that are proportional to *R*_*p*_ and *R*_*d*_, respectively, describe the rate of change of average occupancy of site (*i, j*) due to proliferation and death. An important point regarding this discrete conservation statement is that we are explicitly assuming the occupancy status of lattice sites are independent since we interpret products of lattice site occupancy status as net transition probabilities [49], which is often referred to as the mean-field approximation. Of course, in reality the occupancy status of lattice sites are not independent and we will see some specific examples of this later [1].

To proceed to the continuum limit we identify the quantity ⟨*c*_*i,j*_⟩ with a smooth function *C*(*x, y, t*), and then each term in Equation (1) is expanded in truncated Taylor series about the point (*x, y*) = (*i*Δ, *j*Δ), and terms of 𝒪(Δ^3^) are neglected. We consider the limit as Δ → 0, giving

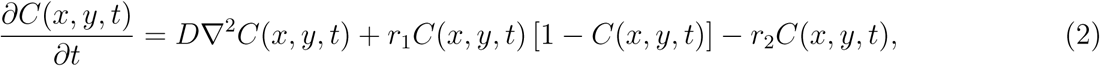

where 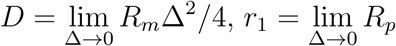 and 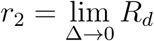. This continuum-limit partial differential equation (PDE) model is a two-dimensional analogue of the well-known Fisher-Kolmogorov model [11,26,39] with an additional linear death term.

In this work we will focus on two different problems inspired by important cell biology experiments. The first problem we consider is inspired by the geometry and initial condition used in a cell proliferation assay [50]. For this problem we consider a square-shaped domain with *I* = *J*. These simulations are initiated by placing a fixed number of agents on the lattice so that each lattice site is occupied with constant probability *C*(0) ∈ [0, 1]. This initial condition means that the density of agents is independent of position so that ∇*C* = **0** and the PDE continuum limit description for *C*(*x, y, t*) simplifies to an ordinary differential equation (ODE) for *C*(*t*),

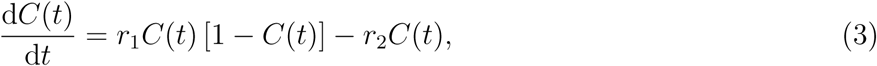

with solution

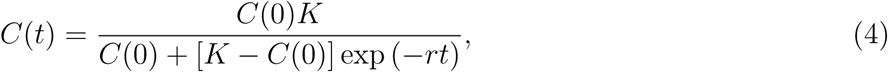

where *K* = (*r*_1_ − *r*_2_)*/r*_1_ and *r* = *r*_1_ − *r*_2_. To compare discrete simulations with the continuum limit in this case we consider a simulation where *N* (0) = ⌊*C*(0)(*I* + 1)(*J* + 1)⌋ agents are distributed randomly on the lattice, taking care to ensure that each lattice site can be occupied by, at most, a single agent. As the simulation proceeds we record the total number of agents on the lattice *N* (*t*), and we convert this time series of agent number to a time series of agent densities by *C*(*t*) = *N* (*t*)*/*(*I* + 1)(*J* + 1). In addition to simply visualising agent locations on the lattice (Figure 1a) we quantitatively summarise the simulation by recording *N* (*t*) at unit time intervals, *t* = 1, 2, 3, …, and we plot the density, which can be compared with the solution of the continuum-limit model (4).

The second problem we consider is inspired by the geometry and initial condition used in a barrier assay [49]. For this problem we consider a narrow rectangular-shaped domain where the horizontal length of the lattice is much larger that the vertical height of the lattice, *I* ≫ *J*. Discrete simulations are initiated by populating lattice sites in the *i*th column with a constant density *C*_*i*_(0) ∈ [0, 1]. This initial condition means that the density of agents is independent of vertical position on the lattice, and the PDE continuum limit description for *C*(*x, y, t*) simplifies to a PDE for *C*(*x, t*) [49], given by

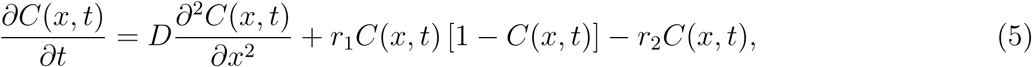

which we solve numerically using an explicit finite different approximation with variable time stepping and automatic truncation error control, as outlined in Section S1 of the Supplementary Material. As mentioned, discrete simulations are initiated by populating lattice sites in each column with a constant probability *C*_*i*_(0) for *i* = 0, 1, 2, …, *I*. The discrete model is allowed to evolve in time and we quantify the outcome of the simulation by counting the number of agents in each column *N*_*i*_(*t*) ∈ [0, *J*] for *i* = 0, 1, 2, …, *I*. These counts are converted into a measure of agent density per column *C*_*i*_(*t*) = *N*_*i*_(*t*)*/J* ∈ [0, 1] for *i* = 0, 1, 2, …, *I*. In addition to simply visualising agent locations on the lattice (Figure 1b) we summarise the simulation output by plotting *C*_*i*_(*t*) as a function of position *x*, recalling that *x* = *i*Δ.

### 2.2 Comparing averaged data from the discrete model and the solution of the continuum limit models

Results in Figure 2(a)–(c) summarise a series of continuum-discrete comparisons for a cell proliferation assay, each with *C*(0) = 0.1, *R*_*m*_ = 1 and *R*_*d*_ = 0.0. The results in Figure 2(a)–(c) compare data from the discrete model with Equation (4) for three scenarios with increasing *R*_*p*_ and we see that the solution of the continuum model accurately approximates data from the discrete model only when *R*_*p*_ is sufficiently small. As *R*_*p*_ increases, the continuum model predicts that *C*(*t*) increases faster than observed using the discrete model. Results in Figure 2(d)–(f) compare data from discrete simulations with Equation (4) for *C*(0) = 0.1, *R*_*m*_ = 1, and *R*_*p*_*/R*_*d*_ = 2. The main difference between the results in Figure 2(a)–(c) and those in Figure 2(d)–(f) is the long-time density of agents on the lattice since we have 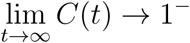 for *R*_*d*_ = 0, whereas we have 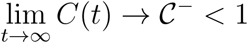 when *R*_*d*_ *>* 0. Again, when *R*_*p*_ and *R*_*d*_ are sufficiently small we see that the solution of the continuum model is accurate, whereas increasing *R*_*p*_ and *R*_*d*_ means that the solution of the continuum model fails to match the data from the discrete model. Comparing results in Figure 2(a)–(c) with those in Figure 2(d)–(f) shows that the inclusion of agent death leads to a greater discrepancy between the solution of the continuum model and data from the stochastic model [1].

**Figure 2:**
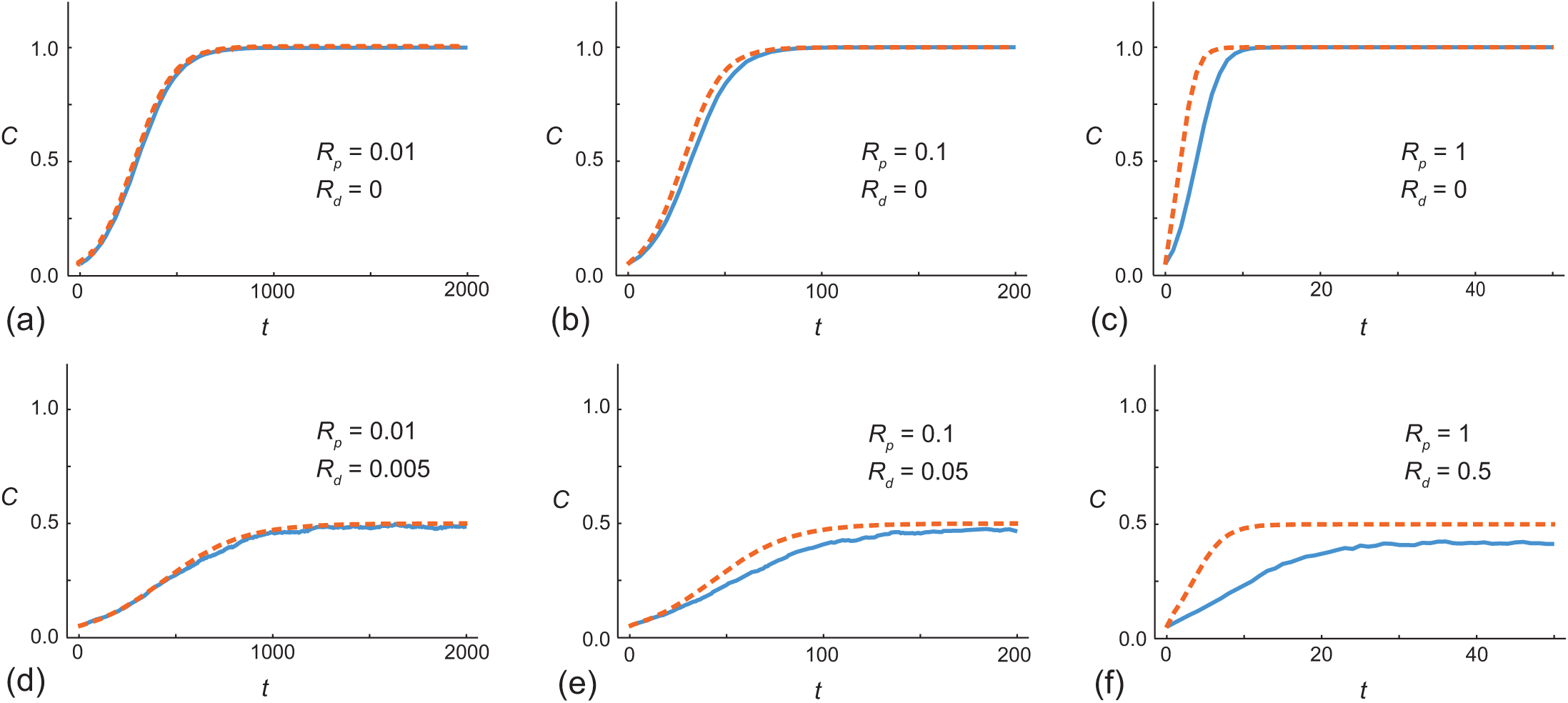
(a)-(c) Evolution of *C*(*t*) for a movement-proliferation model without death. (d)–(f) Evolution of *C*(*t*) for combined movement-proliferation-death. Each figure shows stochastic data (solid blue), the solution of the mean-field continuum model (dashed orange). Stochastic simulations are performed on a lattice with *I* = *J* = 100, with *C*(0) = 0.1.

Note that, in Figure 2(a) we perform simulations over a relatively long time scale, *t* ≤ 2000 so that we can clearly observe the impact of agent proliferation, which takes place at a relatively small rate. In Figure 2(b)–(c) we perform simulations over smaller time scales, *t* ≤ 200 and *t* ≤ 40, respectively. This difference in the choice of time scale reflects the fact that higher rates of proliferation lead to a more rapid increase in *C*(*t*).

Results in Figure 3(a) compare *C*_*i*_(*t*) from the discrete model with the solution of Equation (5). For this problem we have *C*_*i*_(0) = 1.0 for *i* ∈ [90, 110] and *C*_*i*_(0) = 0 elsewhere, with *R*_*m*_ = 1, *R*_*p*_ = 0.01 and *R*_*d*_ = 0. The comparison between the discrete data and the solution of the continuum model is excellent, showing that the population expands outwards, leading to the formation of moving fronts that eventually spread across the whole lattice. The continuum-discrete comparison in Figure 3(b) is for precisely the same problem except that here *R*_*p*_ = 1. Again, we see that the population spreads out across the domain. However, the continuum model does not provide an accurate prediction of the stochastic data in this case. Instead, the continuum model predicts that the population spreads across the domain faster than the discrete model. Again we note that the timescale of the simulations with smaller *R*_*p*_ is longer than the timescale of the simulations with larger *R*_*p*_ so that we allow sufficient time to see the effect of the proliferation mechanism.

**Figure 3:**
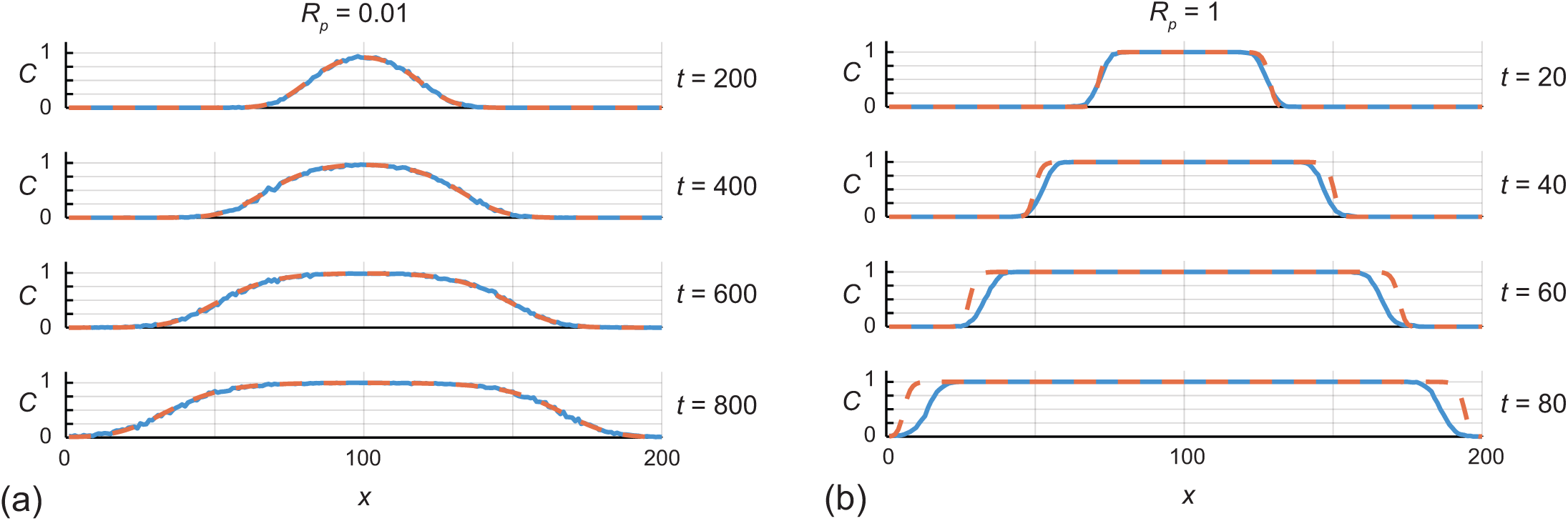
Evolution of *C*(*x, t*) for a movement-proliferation model without death. (a) Evolution of *C*(*x, t*) with *R*_*m*_ = 1 and *R*_*p*_ = 0.01. (b) Evolution of *C*(*x, t*) with *R*_*m*_ = *R*_*p*_ = 1. Each figure shows stochastic data (solid blue), the solution of the mean-field continuum model (dashed orange). All simulations are performed on a lattice with *I* = 200 and *J* = 500, and the initial condition is *C*_*i*_(0) = 1 for *i* ∈ [90, 110] and *C*_*i*_(0) = 0 elsewhere.

Now that we have presented both the discrete and coarse-grained continuum models for the proliferation assay and barrier assays in Figures 2–3, it is insightful to explore the computational expense of both approaches. Stochastic simulation results in Figure 2(a) require approximately 7 seconds of computation time, whereas the coarse-grained model requires approximately 0.00003 seconds to compute, which is six orders of magnitude faster. Of course, since the discrete model is stochastic the computation time varies slightly each time we simulate the same identically-prepared initial condition. However, regardless of this variability, the coarse-grained model is leads to a computational speed up of six orders of magnitude. The coarse-grained model for the proliferation assay is an ODE that has an exact solution so it is not surprising that the computation time is extremely fast. Stochastic simulation results in Figure 3(a) for the barrier assay requires approximately 240 seconds of computation time whereas the coarse-grained model requires approximately 0.06 seconds to compute, which is five orders of magnitude faster. Here, the coarse-grained model for the barrier assay is a PDE that we solve numerically, but we still find that the numerical solution of this PDE model is significantly faster than performing stochastic simulations. All computations are performed on a standard laptop using with an Intel i7-1185G7 CPU (3.00GHz).

At this point it is useful to discuss notation. So far we use *C*(*t*) to denote to the density of agents in a proliferation assay at time *t*, and we use this same notation regardless of whether we consider a stochastic simulation or the solution of the continuum-limit model, Equation (4). Similarly we use *C*(*x, t*) to denote to the density of agents at position *x* and at time *t* in a barrier assay, as given by the solution of Equation (5), or we use *C*_*i*_(*t*) to denote the density of agents in the *i*th column at time *t* in a stochastic simulation of a barrier assay. To address the question of parameter inference, we will be more explicit by using *C*(*t* | *θ*) to denote the density of agents in a proliferation assay at time *t* with parameters *θ*. Similarly, we use *C*(*x, t* | *θ*) and *C*_*i*_(*t* | *θ*) to note the density of agents in a barrier assay for the continuum-limit model and stochastic model, respectively.

### 2.3 Reliable predictions incorporating model error: Proliferation assay

We now describe an approach for incorporating model error between the discrete and continuum models. Throughout this section we will refer to the fine (discrete, stochastic) model as *F* (*θ*), and the coarse continuum-limit model as *g*(*θ*), where *θ* is the vector of parameters of length 𝒩_*p*_. We have used a capital letter *F* for the fine model to indicate that this is stochastic and we denote a specific realisation of *F* for some *θ* by *f* (*θ*). Our approach has two core components: (i) an evaluation of the model error at a finite number of locations; and (ii) the construction of a statistical meta-model of this error on the basis of these samples. We describe each of these next.

#### 2.3.1 Model error samples

The first core component of our approach to statistically characterising the model error begins by evaluating the model error at a finite number of sample locations. A maximum computational budget of fine-model evaluations usually bounds this number of evaluations. Thus we:

1. Sample vector realisations *θ*^*i*^ for *i* = 1, 2, 3, …, 𝒩_*d*_, from a particular design distribution that we will specify later.
2. For each *θ*^*i*^, *i* = 1, 2, 3, …, 𝒩_*d*_, simulate the fine model to give *f* (*θ*^*i*^) ∼ *F* (*θ*^*i*^), and solve the coarse model to give *g*(*θ*^*i*^).
3. Define *ε*^*i*^ = *f* (*θ*^*i*^) − *g*(*θ*^*i*^), which is a vector of length 𝒩_*t*_, for each *i* = 1, 2, 3, …, 𝒩_*d*_. Here, *f* (*θ*) denotes the prediction of the fine model and *g*(*θ*) denotes the prediction of the coarse model. Each model prediction is a time series at time points 1, 2, 3, …, 𝒩_*t*_,
4. Record (*ε, θ*)^*i*^ = (*ε*^*i*^, *θ*^*i*^) for *i* = 1, 2, 3, …, 𝒩_*d*_.

#### 2.3.2 Statistical meta-model of the model error

The next core component is to construct a statistical (meta-)model of *ε* from the 𝒩_*d*_ evaluations. As discussed earlier, in the traditional literature on the design of computer experiments, the simulation model to approximate (i.e. the fine-scale model here) is usually considered deterministic, and this raises the question of the most appropriate place to employ probability modelling [25, 41, 47]. In a Bayesian approach, this is no real conceptual difficulty as probability reflects personal uncertainty; a frequentist approach is perhaps less obvious, but Owen [25, 41] developed a method in which the only actual randomness enters via the (known) design distribution with target criteria defined with respect to this sampling distribution. However, in the present case, the fine-scale model is stochastic. These conceptual issues are thus less critical: randomness can enter both via the design distribution and the variability of the stochastic fine-scale model. Our approach here is to allow for flexibility in the choice of design distribution – this can be chosen on intuitive grounds (e.g. similarly to a prior distribution), according to (parameter) *space-filling* criteria [12, 25, 41], or according to some other formal experimental design criterion. For the main results, we will adopt a simple uniform sampling design. However, we also consider how these results are impacted by choosing truncated multivariate normal and Latin hypercube designs in Appendices B–C, respectively.

Regardless of design distribution chosen, this stage involves a choice of which class of statistical meta-model to use for the resulting model error. Numerous flexible models, such as Gaussian processes, neural networks, or support vector machines, are often considered in surrogate modelling [12, 15]. Here, however, the availability of a reasonably accurate approximate coarse model with a clear interpretation means we only need to model the *error* between these two models, which in our case is the stochastic model and the continuum-limit model. We likely cannot expect this error term to be mean zero, reflecting potential bias between the models. We may also quite reasonably expect it to have some parameter dependence: we know that the coarse model is a better approximation in some parameter regimes than others as illustrated in Figures 2–3. However, it is perhaps plausible that the mean error may only depend (approximately) linearly on the parameters for a good coarse model. Thus we adopt the simple approach of constructing a *linear regression meta-model for the model error term*. This linear relationship need not hold exactly. It is true that the expressions for the conditional mean and covariance we will derive under this approximation are exact when the joint distribution of (*ε, θ*) is multivariate normal [7]. More importantly, however, the expressions can also be interpreted in terms of ‘best linear approximations’ (or linear projections of the true model) even when the true conditional expectation is not exactly linear (though this approximation is of course more relevant the more plausible it is) [7, 59].

We thus next construct a multivariate linear least squares approximation. These are elementary results from multivariate statistical modelling [7, 10], but we find it useful to give the steps explicitly to facilitate understanding and implementation. Starting from the samples obtained from the above procedure, (*ε*^*i*^, *θ*^*i*^) for *i* = 1, 2, 3, …, 𝒩_*d*_, we arrange and store the data in the following block format:

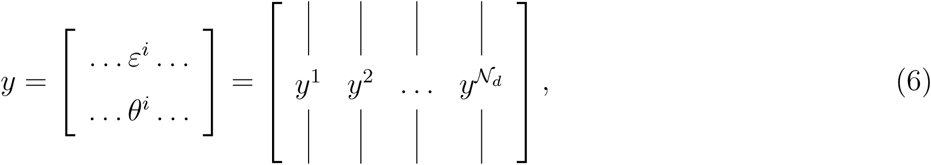

for *i* = 1, 2, 3, …, 𝒩_*d*_, where 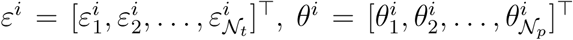, and *y*^*i*^ = [*ε*^*i*⊤^, *θ*^*i*⊤^]^⊤^, with dimensions as defined previously. Note that, in keeping with statistical terminology, we do not use boldface for vectors: for example, *ε*^*i*^ is the *i*th sample of an 𝒩_*t*_-dimensional vector with components 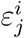.

With this data we calculate the (empirical) row mean of *y* over the 𝒩_*d*_ samples,

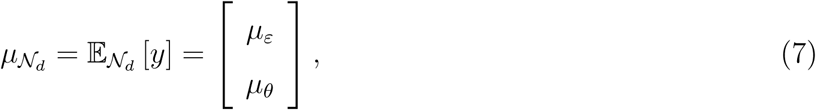

where *μ*_*ε*_ is a vector of length 𝒩_*t*_ and *μ*_*θ*_ is a vector of length 𝒩_*p*_, and then calculate the block structured (empirical) covariance matrix

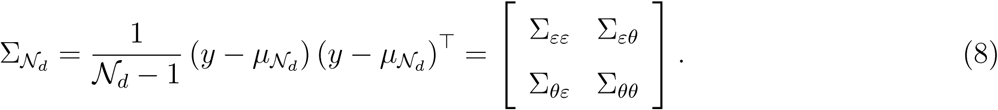

The Σ_*εε*_ block is of size 𝒩_*t*_ × 𝒩_*t*_, the Σ_*θθ*_ block is of size 𝒩_*p*_ × 𝒩_*p*_, and the 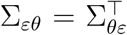 block is of size 𝒩_*t*_ × 𝒩_*p*_.

With this data we compute the (linear approximation to the) conditional mean and conditional covariance of *ε* given *θ*

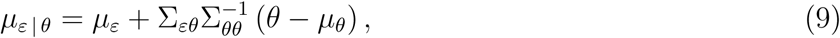

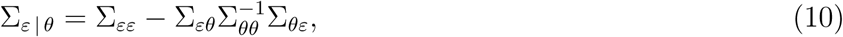

where the term 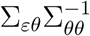 corresponds to the multivariate linear regression coefficients [7]. As mentioned above, Equations (9)-(10) can be considered as valid ‘linear least squares’ approximations regardless of the actual distributions of *ε* and *θ*, and are the finite-sample analogues of the population (infinite data) linear approximation model [59]. However, we can further strengthen our working assumptions based on these two moments and (for now) work with the simple, stronger statistical model for the correction term:

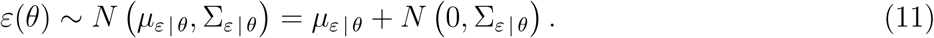

With this framework we now approximate the prediction of the fine model in terms of the coarse model and the discrepancy as

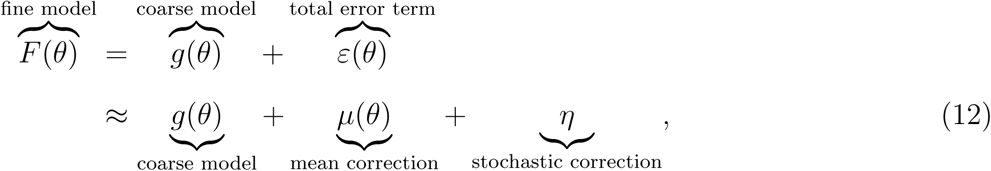

where *μ*(*θ*) = *μ*_*ε* | *θ*_ is the parameter-dependent, deterministic (conditional on a design realisation) linear approximation to the conditional mean of the model error and *η* ∼ *N* (0, Σ_*ε* | *θ*_) is a parameterindependent (as the expression for the conditional covariance is independent of *θ*), mean-zero stochastic error representing the remaining components of model error.

We now demonstrate how to make reliable predictions using this framework. First, in Figure 4, we work with the stochastic model of a cell proliferation assay with *C*(0 | *θ*) = 0.1 and *R*_*m*_ = 1.0 both held fixed. We sample 𝒩_*d*_ = 500 pairs of [*R*_*p*_, *R*_*d*_] uniformly, at random, in the region *R*_*p*_ ∈ [0.8, 1.2] and *R*_*d*_ ∈ [0.0, 0.4] and consider simulations for *t* ∈ [0, 30]. As we showed in Figure 2, the solution of the continuum model in this region of parameter space is a relatively poor approximation of the stochastic simulation data. Results in Figure 4(a) illustrate the 500 uniformly-sampled pairs of [*R*_*p*_, *R*_*d*_], and results in Figure 4(b)–(d) show *C*(*t* | *θ*) for three particular choices of parameters: (i) [*R*_*p*_, *R*_*d*_] = [0.8, 0.2]; (ii) [*R*_*p*_, *R*_*d*_] = [0.9, 0.3]; and, (iii) [*R*_*p*_, *R*_*d*_] = [0.7, 0.1]. In each case we see that the dynamics of the relatively expensive fine (stochastic) model (solid blue) are poorly predicted by the computationally efficient coarse (continuum) model (dashed orange). Fortunately, simply adding the mean correction term, *μ*(*θ*), to the solution of the coarse (continuum) model, *g*(*θ*), we obtain a corrected coarse model (dotted red) that combines the best of both approaches – solutions are extremely fast to compute and the model accurately captures data from the expensive fine (stochastic) model. Results in Figure 4 compare the solutions of the fine (stochastic), coarse (continuum) and corrected coarse models for three particular choices of [*R*_*p*_, *R*_*d*_]. Additional comparisons of the three models for other choices of [*R*_*p*_, *R*_*d*_] in the region sampled in Figure 4(a) lead to similar results (not shown).

**Figure 4:**
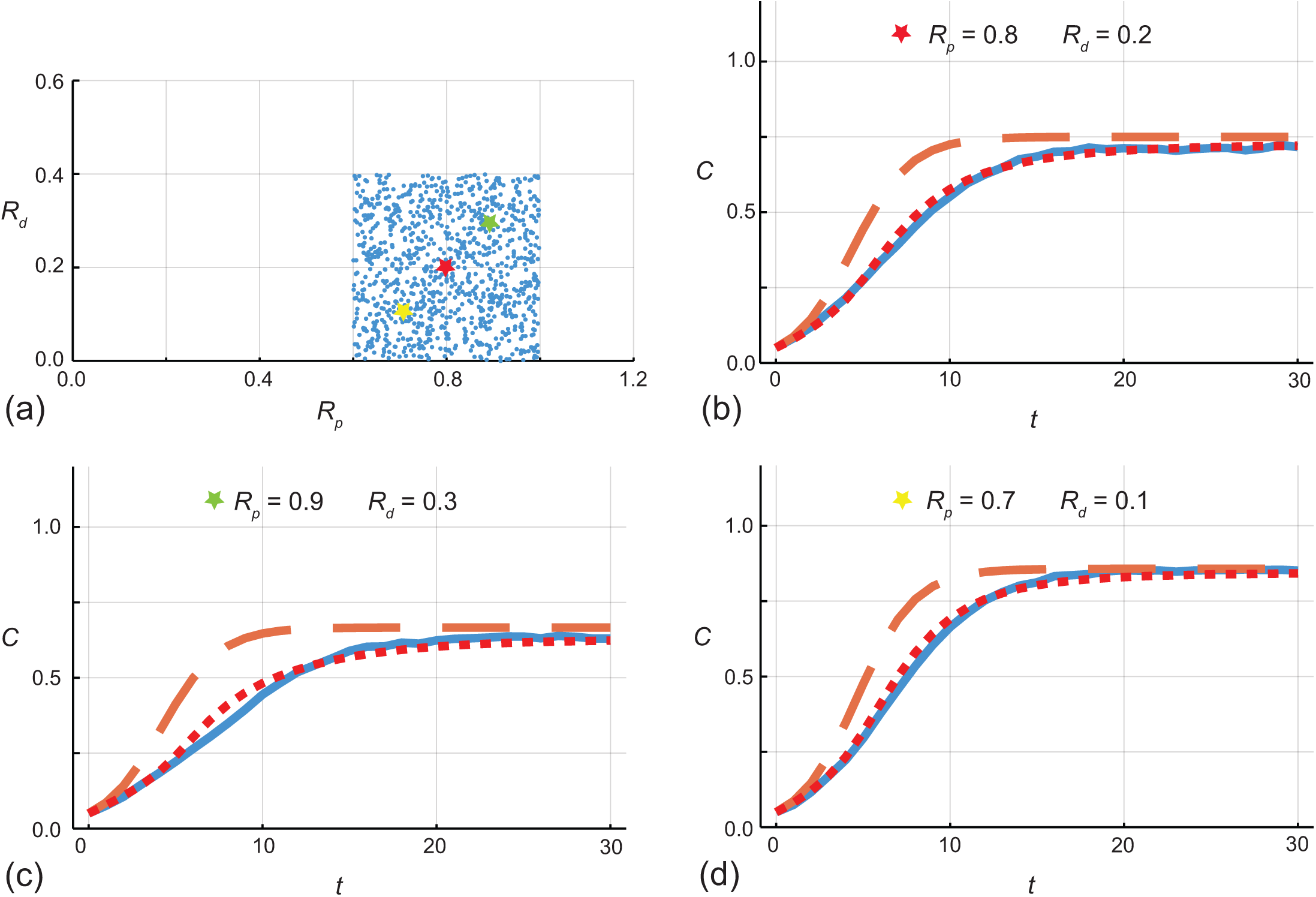
(a) 𝒩_*d*_ = 500 uniform samples of pairs of [*R*_*p*_, *R*_*d*_] within a square-shaped region in the interval *R*_*p*_ ∈ [0.0, 1.0] and *R*_*d*_ ∈ [0.0, 0.4] are used to construct the error model. (b)–(d) Comparison of *C*(*t* | *θ*) for the fine (stochastic) model (solid blue), the coarse (continuum) model (dashed orange) and the coarse model plus mean correction term (dotted red) for [*R*_*p*_, *R*_*d*_] = [0.8, 0.2], [0.9, 0.3] and [0.7, 0.1], respectively. Stochastic simulations are performed on a lattice with *I* = *J* = 100, with *C*(0 | *θ*) = 0.1 and *R*_*m*_ = 1. Stochastic simulation data are recorded at *t* = 1, 2, …, 29, 30, giving 𝒩_*t*_ = 30.

Each evaluation of the corrected coarse model in Figure 4 is deterministic, given a fixed set of design points from the first stage, in the sense that we have taken the correction term to be deterministic by setting *η* = 0 in Equation (12) for convenience. Of course, it is straightforward to include variability into the correction term simply by evaluating and incorporating *η* as given in Equation (12). Results in Figure 5 show data from the stochastic model for the same three choices of parameters from Figure 4. For each parameter choice we superimpose 50 realisations of the corrected coarse model that includes the stochastic term, *η*, in Equation (12). This simple visual comparison of data from the full stochastic model with several realisations of the corrected model, this time including the *η* term, illustrates the versatility of this approach. In this work we will consider both approaches by setting *η* = 0 or incorporating *η* where convenient.

**Figure 5:**
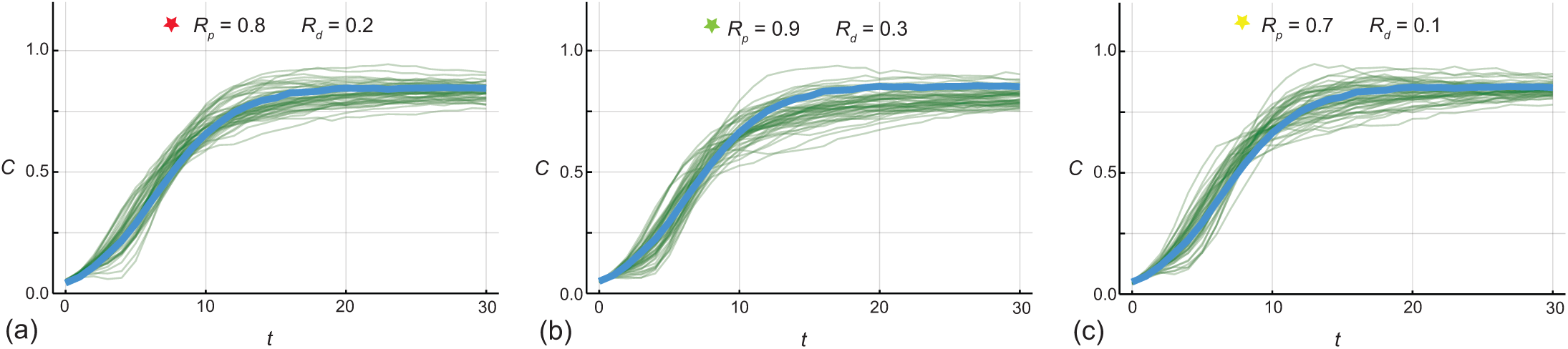
(a)–(c) Comparison of *C*(*t* | *θ*) from the stochastic model (solid blue) for [*R*_*m*_, *R*_*p*_, *R*_*d*_] = [0.8, 0.2], [0.9, 0.3], and [0.7, 0.1], respectively. In each subfigure we superimpose the solution of the continuum model plus the mean and 50 realisations of the stochastic correction terms (thin green opaque). Stochastic simulation data are recorded at *t* = 1, 2, …, 29, 30, giving 𝒩_*t*_ = 30.

Now we have demonstrated how to incorporate both the mean and stochastic correction terms to make reliable predictions using a (potentially) inaccurate continuum limit, we will address to the question of using the correction terms to improve the more challenging computational task of parameter inference.

### 2.4 Reliable inference incorporating model error: Proliferation assay

We will consider parameter inference using data generated by the stochastic model, denoted by a vector *C*^o^ with components corresponding to 𝒩_*t*_ discrete times, *t*_*i*_, for *i* = 1, 2, 3, …, 𝒩_*t*_. Observed data are denoted with a superscript ‘o’ to distinguish these noisy observations from the various modelled prediction, *C*(*t* | *θ*), which we will obtain using several methods outlined below. For the proliferation assay we will evaluate *C*(*t* | *θ*) at *t* = 1, 2, 3, …, 𝒩_*t*_. For completeness we include additional additive and normally distributed measurement error with zero mean and constant variance, i.e. *C*^o^ | *θ* ∼ *N* (*C*(*t* | *θ*), *σ*^2^*I*) where *I* is the 𝒩_*t*_ × 𝒩_*t*_ identity. While in principle the variance associated with the observation error can be estimated along with the other components of *θ* [52], for simplicity we will pre-specify *σ*^2^ [17].

In this work we take a likelihood-based approach to parameter inference and uncertainty quantification. For a time series of observations together with the standard noise assumptions outlined above, the log-likelihood function is

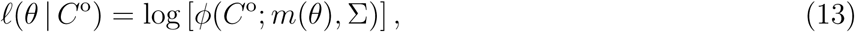

where the observations, *C*^o^, are taken from a single realisation of the stochastic model and *ϕ*(*x*; *m*(*θ*), Σ) denotes a multivariate normal probability density function with mean *m*(*θ*) and covariance Σ, as above. For clarity, recall that *m*(*θ*) is a vector of length 𝒩_*t*_ and Σ is a covariance matrix of size 𝒩_*t*_ × 𝒩_*t*_. We have written the likelihood function in Equation (13) in terms of a generic mean vector and covariance matrix, however, as we will use particular forms as follows. When we work with the mean correction we will use *m*(*θ*) = *C*(*t* | *θ*) + *μ*(*θ*) and Σ = *σ*^2^*I*, where *C*(*t* | *θ*) is the solution of the coarse model. In contrast, when we work with the mean and stochastic correction we take *m*(*θ*) = *C*(*t* | *θ*) + *μ*(*θ*) and Σ = Σ_*ε* | *θ*_ + *σ*^2^*I*, where again *C*(*t* | *θ*) is the solution of the coarse model. In summary, in this framework we compute 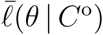in three ways, using: (i) the coarse model without any correction; (ii) the coarse model with the mean correction; and, (iii) the coarse model with both the mean and stochastic correction. We will demonstrate the different ways to compute and interpret 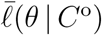later in this section.

Applying maximum likelihood estimation (MLE) gives a best fit set of parameters, 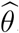. The MLE is defined as

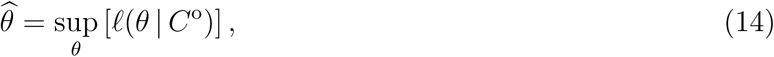

which we compute numerically using NLopt in Julia [19] with the BOBYQA optimization algorithm and user-specified bound constraints [44].

In this work we use a profile likelihood-based approach to explore uncertainty quantification and practical identifiability [43, 45, 52, 57]. In all cases we work with a normalised log-likelihood function

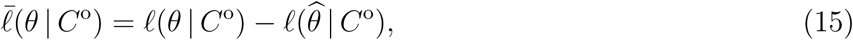

which we interpret as a function of *θ* for fixed data *C*^o^. We choose to work with the normalised log-likelihood function because it allows us to visually and quantitative compare different likelihood functions more easily. Assuming the parameter vector *θ* can be partitioned into an *interest* parameter *ψ* and a *nuisance* parameter *λ*, we write *θ* = (*ψ, λ*). Given a set of data, *C*^o^, the profile log-likelihood for the interest parameter *ψ* can be written as

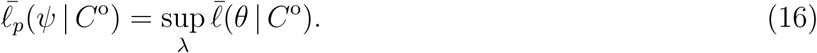

In Equation (16), *λ* is optimised out for each value of *ψ* to implicitly define a function *λ*^*^(*ψ*) of optimal values of *λ* for each value of *ψ* [52]. In this work we estimate *λ*^*^(*ψ*) by evaluating this function across a grid of values of *ψ*. For example, in the simulations of a proliferation assay where we treat *C*(0 | *θ*) and *R*_*m*_ = 1 as fixed, we have *θ* = (*R*_*p*_, *R*_*d*_). If we wish to profile the proliferation rate then we have *ψ* = *R*_*p*_ and *λ* = *R*_*d*_ so that

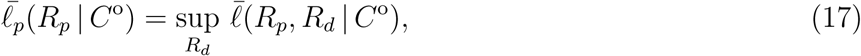

which again we compute numerically using NLopt in Julia [19] with the BOBYQA optimization algorithm with user-specified bound constraints [44]. For each value of the interest parameter, taken over a sufficiently fine grid, the nuisance parameter is optimised out and the previous optimal value can be used as the starting estimate for the next optimisation problem as we move along the fine grid. In this work we use uniformly spaced grids of 100 points defined on problem-specific intervals that are reported in the plots of the various profile likelihoods.

To demonstrate these ideas we consider stochastic data in Figure 4(b) for *R*_*p*_ = 0.8 and *R*_*d*_ = 0.2, with data generated from the stochastic model, *C*^o^ = *N* (*t*)*/*(*I* + 1)(*J* + 1) for *t* ∈ [0, 30], and *C*^o^ is recorded at *t* = 1, 2, 3, …, 30, giving 𝒩_*t*_ = 30. With this data we compute 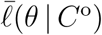 over the region *R*_*p*_ ∈ [0.6, 1.0] and *R*_*d*_ ∈ [0.0, 0.40] using a uniform mesh on the parameter space. At each point on the mesh we compute 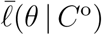 in three ways, as described previously. Heat maps of 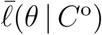 are shown in Figure 6 with a red disc superimposed at the parameter values used to generate the stochastic data, which we will informally refer to as the *true value*. Each heat map in Figure 6 is superimposed with contours showing 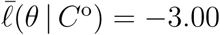, which defines an approximate 95% confidence region [46, 52]. Without incorporating any correction term we have poor coverage since the true value is (consistently) excluded from the 95% confidence region, whereas incorporating either the mean correction term only or the mean plus stochastic correction terms leads to the true value being contained within the 95% confidence region.

**Figure 6:**
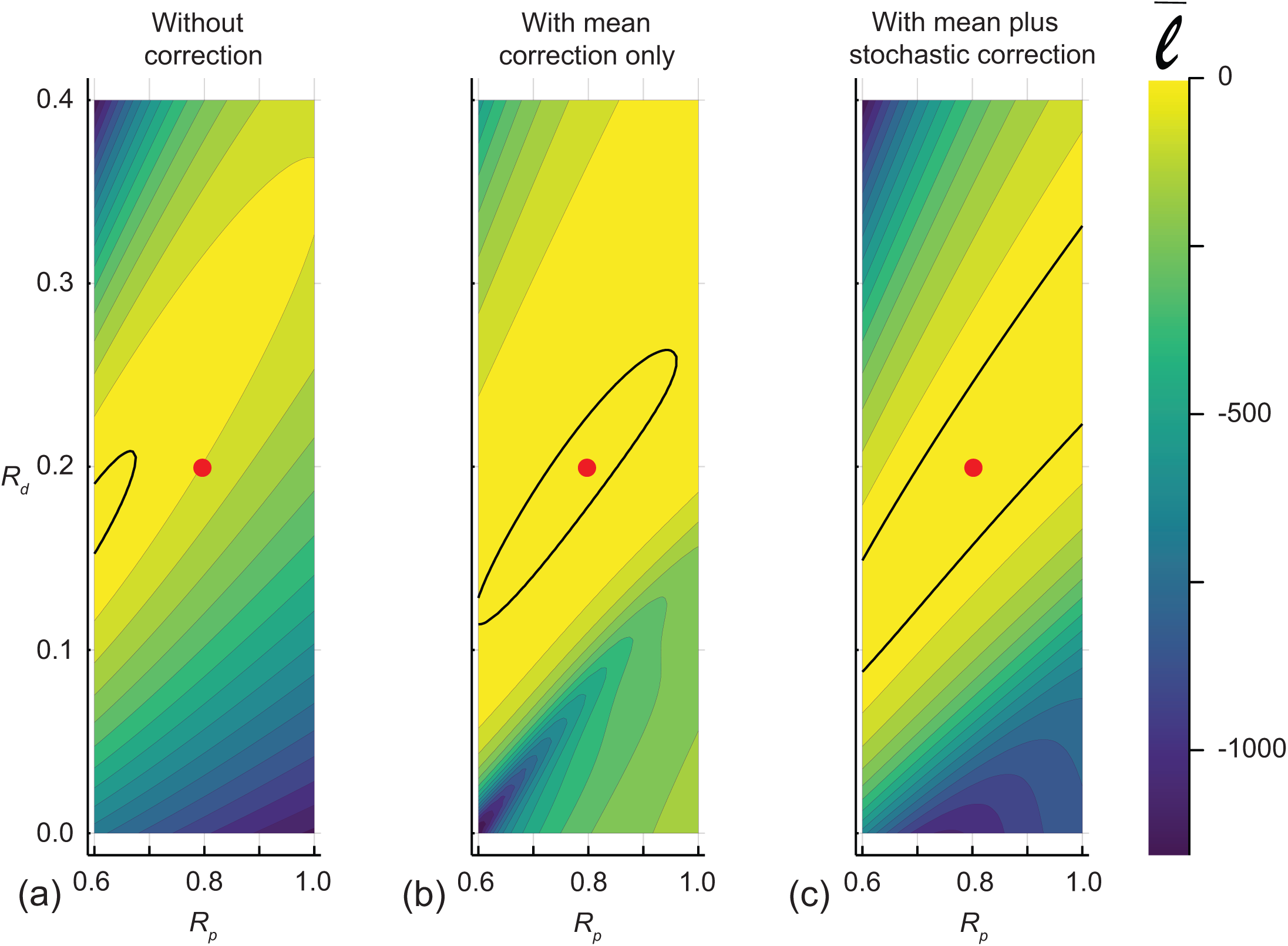
(a)–(c) show the normalised log likelihood as a function of [*R*_*p*_, *R*_*d*_] without any correction, with the mean correction only, and with the mean plus stochastic correction, respectively. Each heat map is constructed by evaluating the likelihood across a 40 × 40 uniform discretisation of the parameter space. The outline of the region where 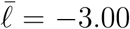defines the approximate 95% confidence region and the location of the values used to generate the data, [*R*_*p*_, *R*_*d*_] = [0.8, 0.2], is superimposed on each plot. All calculations were carried out with *σ*^2^ = 0.05.

To further explore inference and uncertainty quantification for the likelihood functions in Figure 6 we construct several profile likelihoods, summarised in Figure 7. Figure 7(a) shows the univariate profiles for *R*_*p*_ and *R*_*d*_, without either of the correction terms, where we see that the MLE underestimates the true values of both *R*_*p*_ and *R*_*d*_. In this work, each univariate profile is superimposed with a horizontal dashed line at 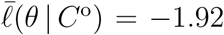 which defines an approximate 95% confidence interval on the univariate profile likelihood function [46]. We see that without incorporating any correction, the apparent 95% confidence intervals do not contain the true values. Results in Figure 7(b)–(c) show the univariate profiles with the mean correction and with the mean plus stochastic correction, respectively. Comparing the profiles in Figure 7(a) and (b) we see that incorporating the mean correction leads to profiles that contain the true value, and the MLE is quite close to the true values. Comparing the profiles in Figure 7(b) and (c) shows that incorporating the stochastic correction widens the profiles, and again the MLE is quite close to the true value. The widening of the profiles in Figure 7(c) indicates that we have increased uncertainty associated with the additional correction term.

**Figure 7:**
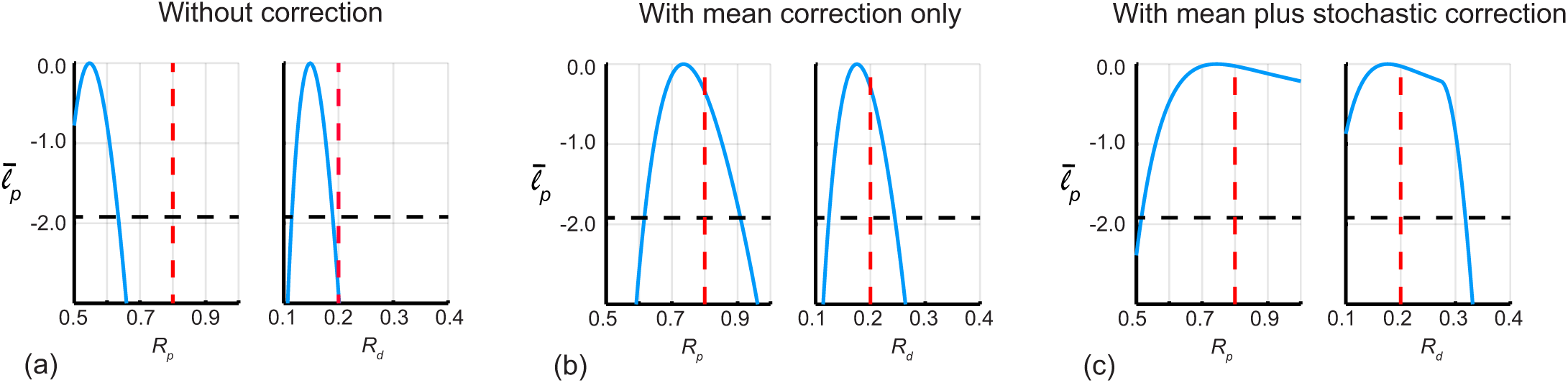
(a)–(c) univariate profile likelihoods, 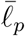, for *R*_*p*_ and *R*_*m*_, as indicated, for the model without any correction, with the mean correction, and with the mean plus stochastic correction, respectively. The horizontal dashed lines indicates the threshold 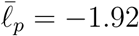 that defines approximate 95% confidence intervals, and the vertical solid red line shows the values used to generate the data. In (a), without the correction, we have 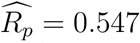, [0.500, 0.634] and 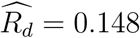, [0.115, 0.189]. In (b), with the mean correction only, we have 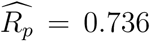, [0.616, 0.908] and 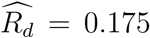, [0.124, 0.244]. In (c), with the mean plus stochastic correction, we have 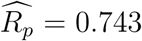, [0.515, 1.000] and 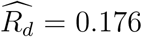, [0.100, 0.318]. All calculations were carried out with *σ*^2^ = 0.05.

### 2.5 Reliable predictions and inference incorporating model error: Barrier assay

All results in Sections 2.3–2.4 are generating using simulations of proliferation assays as reported in Figure 1(a) and Figure 2 where the initial distribution of agents is independent of position. Here, we show that the same concepts extend to the simulations of barrier assays reported in Figure 1(b) and Figure 3 where the density of agents depends upon both position and time. We apply the same ideas except that the data we work with is slightly different. We work with a similar problem to that reported in Figure 3(b) with parameters close to *R*_*m*_ = 0.5, *R*_*p*_ = 2.0 and *R*_*d*_ = 0.0, where the solution of the continuum model, Equation (5), is relatively inaccurate as an approximation to the dynamics of the fine stochastic model. We apply the same approach to correcting the predictions of the coarse (continuum) model as we did in Sections 2.3–2.4 except that we take the simplest approach and record *C*_130_(*t* | *θ*) at *t* = 1, 2, 3, …, 60, giving 𝒩_*t*_ = 60. Note that collecting the density in column 130 corresponds to collecting the time series of density at *x* = *x*_o_ = 130, and for consistency with the results in Sections (2.3)–(2.4) we will refer to *C*_130_(*t* | *θ*) as *C*(*t* | *θ*).

Results in Figure 8(a) show the 𝒩_*d*_ = 300 pairs of [*R*_*m*_, *R*_*p*_] sampled uniformly in the region *R*_*m*_ ∈ [0.3, 0.7] and *R*_*p*_ ∈ [1.8, 2.2]. Figure 8(b)–(d) compares the evolution of *C*(*t* | *θ*) for three particular choices of parameters: (i) [*R*_*m*_, *R*_*p*_] = [0.5, 2.0]; (ii) [*R*_*m*_, *R*_*p*_] = [0.4, 2.1]; and, (iii) [*R*_*m*_, *R*_*p*_] = [0.6, 1.9]. In each case we see that the relatively expensive fine (stochastic) model (solid blue) is poorly predicted by the solution of the computationally efficient coarse (continuum) model (dashed orange). As with the results in Section 2.3, simply adding the mean correction term to the solution of the coarse (continuum) model, we obtain a corrected coarse model (dotted red) that combines the best of both approaches, producing a computationally inexpensive means of accurately predicting data from the fine (stochastic) model. As shown in Figure 8(b)–(c) the correction term leads to some overshoot for some choices of parameters, but this is not always the case for other parameter choices, as illustrated in Figure 8(d). Additional results (not shown) indicate that the overshoot can be minimised by increasing 𝒩_*d*_, but of course this comes with additional computational cost. Instead, here we have deliberately chosen to work with a modest value of 𝒩_*d*_ and illustrate that even a modest choice of 𝒩_*d*_ leads to significant improvements in both model prediction and parameter inference.

**Figure 8:**
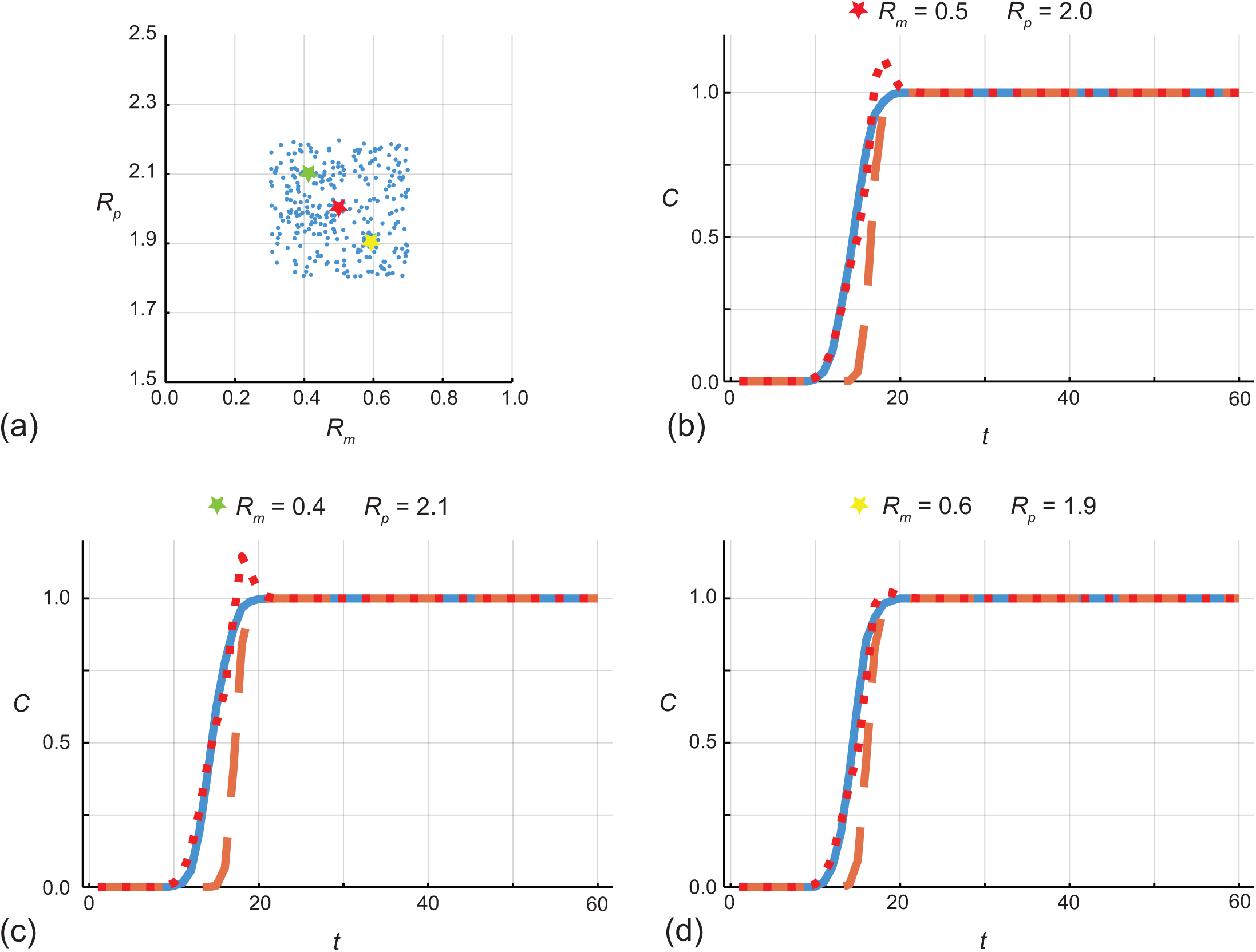
(a) 𝒩_*d*_ = 300 uniform samples of pairs of [*R*_*m*_, *R*_*p*_] within a square-shaped region in the interval *R*_*m*_ ∈ [0.3, 0.7] and *R*_*p*_ ∈ [1.8, 2.2] used to construct the error model. (b)–(d) Comparison of *C*(*t*|, | *θ*) for the fine (stochastic) model (solid blue), the coarse (continuum) model (dashed orange) and the coarse model plus mean correction term (dotted red) for [*R*_*p*_, *R*_*d*_] = [0.5, 2.0], [0.4, 2.1] and [0.6, 1.9], respectively. Stochastic simulations are performed on a lattice with *I* = 200 and *J* = 400 with *C*_*i*_(0 | *θ*) = 1.0 for *i* ∈ [90, 110] and *C*_*i*_(0 | *θ*) = 0 elsewhere. Stochastic simulation data, *C*_130_(*t* | *θ*) = 0, are recorded at *t* = 1, 2, …, 59, 60, giving 𝒩_*t*_ = 60.

While the comparison of models of the barrier assay in Figure 8 deals with the mean correction term only by setting *η* = 0, the comparison in Figure 9 incorporates both the mean and stochastic correction terms. In comparison with Figure 5, the results in Figure 9 are far noisier for the proliferation assay. This difference is due to the fact that the stochastic simulation data we work with for the proliferation assay involves relatively small fluctuations since we work with a relatively large lattice and averages are constructed across the whole lattice. In contrast, stochastic simulation data for the barrier assay involves larger fluctuations since averages are constructed along each column of the lattice only. Indeed, additional results that involve re-computing Figure 5 for a proliferation assay on a smaller lattices leads to larger fluctuations like we see in Figure 9 (not shown).

**Figure 9:**
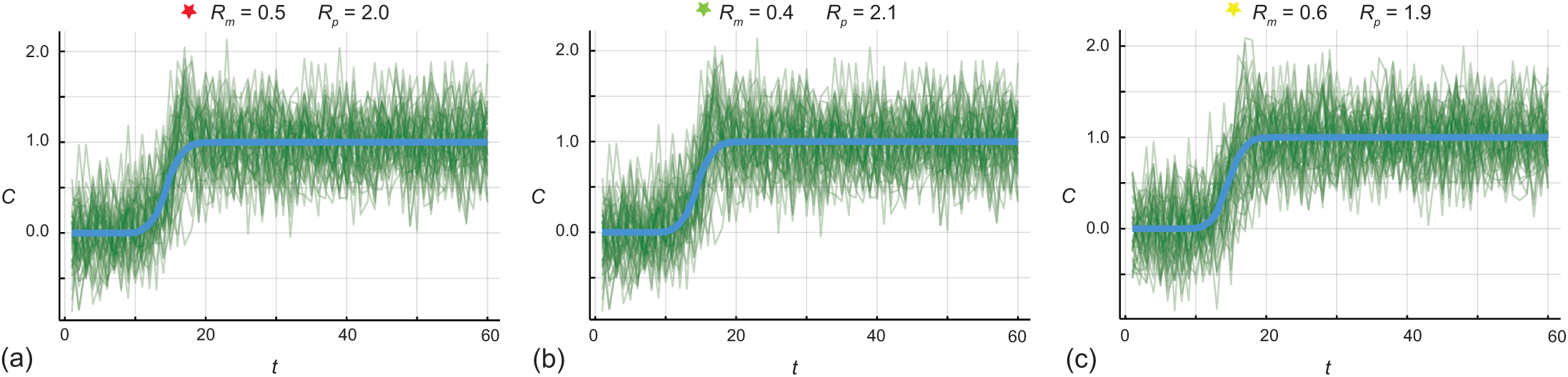
(a)–(c) Comparison of *C*(*t* |,| *θ*) from the stochastic model (solid blue) for [*R*_*m*_, *R*_*p*_] = [0.5, 2.0], [0.4, 2.1], and [0.6, 1.9], respectively. In each subfigure we superimpose the solution of the continuum model plus the mean and 50 realisations of the stochastic correction terms (thin green opaque). Stochastic simulation data, *C*_130_(*t* | *θ*) = 0, are recorded at *t* = 1, 2, …, 59, 60, giving 𝒩_*t*_ = 60.

Turning our attention to inference for the barrier assay simulations we compute 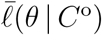 over the region *R*_*m*_ ∈ [0.2, 0.8] and *R*_*p*_ ∈ [1.0, 3.0] using a uniform meshing of the parameter space. At each point on the mesh we compute 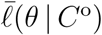in three ways, using: (i) the coarse model without the correction term; (ii) the coarse model with the mean correction; and, (iii) the coarse model with both the mean and stochastic correction. Heat maps of 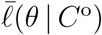 are given in Figure 10 with a red disc superimposed at the true parameter values. Each heat map in Figure 10 is superimposed with contours showing 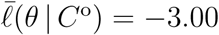, which defines an approximate 95% confidence region. As for the proliferation assay, without the correction term we have poor coverage since the true value is (consistently) excluded from the 95% confidence region, whereas incorporating the mean correction term only or the mean plus stochastic correction terms leads to the true value being contained within the 95% confidence region. Comparing results in Figure 10(b)–(c) shows that incorporating the stochastic correction term broadens the confidence region, which reflects the additional uncertainty included in the likelihood function.

**Figure 10:**
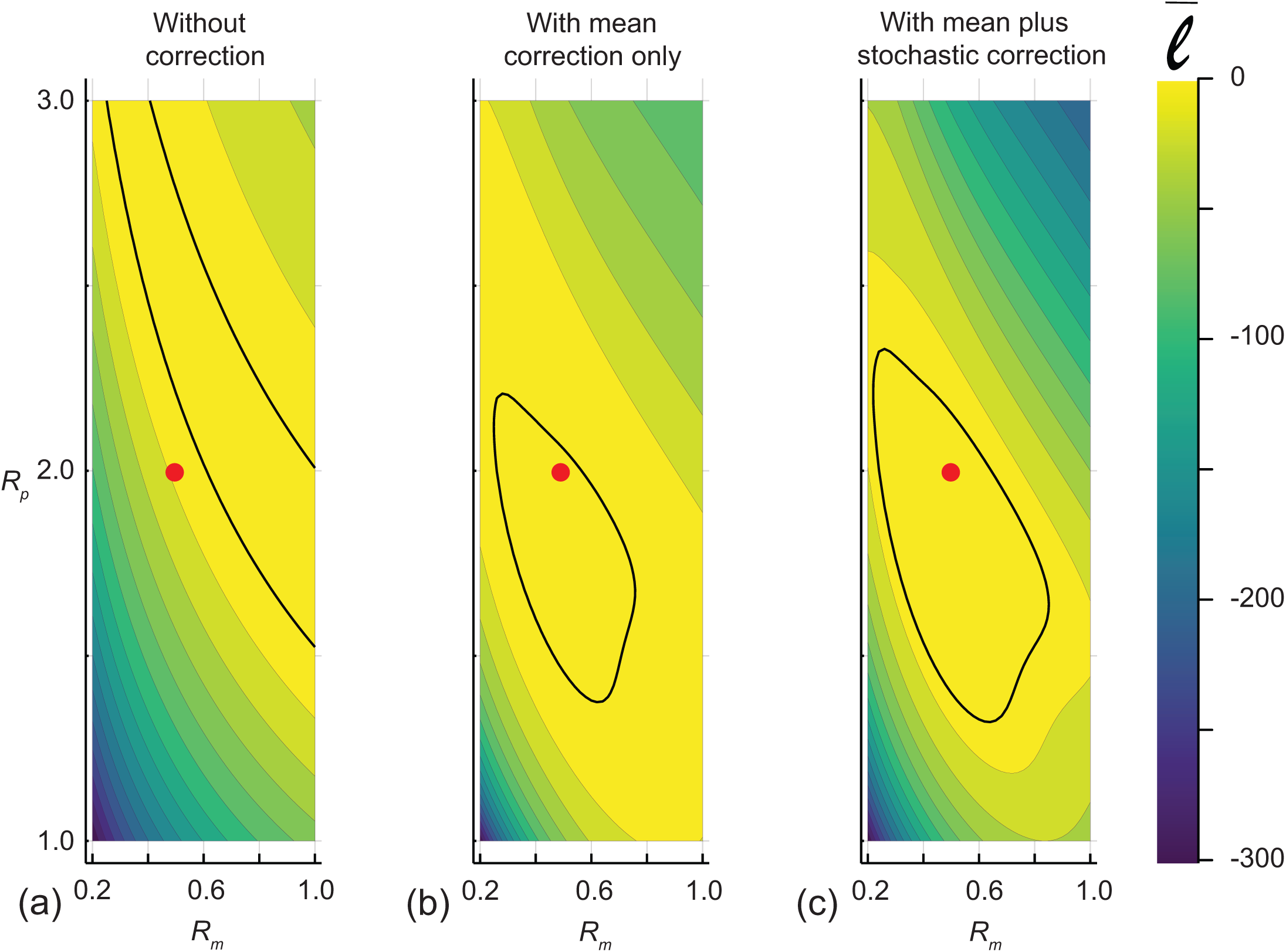
(a)–(c) show the normalised log likelihood as a function of [*R*_*m*_, *R*_*p*_] without any correction, with mean correction only, and with mean plus stochastic correction, respectively. Each heat map is constructed by evaluating the likelihood across a 40 × 100 uniform discretisation of the parameter space. The outline of the region where 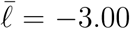 defines the approximate 95% confidence region and location of the values used to generate the data, [*R*_*m*_, *R*_*p*_] = [0.5, 2.0], is superimposed on each plot. All calculations were carried out with *σ*^2^ = 0.05.

The univariate profile likelihoods for the barrier assay problem are given in Figure 11. Figure 11(a) shows the univariate profiles for *R*_*m*_ and *R*_*p*_ for the approximate model without any correction term, showing that the MLE overestimates the true value *R*_*m*_ and underestimates the true value of *R*_*p*_. As with the simulations of the proliferation assay, without incorporating any correction the apparent 95% confidence intervals do not contain the values used to generate the data, which we refer to informally as the *true value*. Figure 10(b)–(c) show the univariate profile likelihoods with the mean correction and with the mean plus stochastic correction, respectively. Similar to the proliferation assay, incorporating the mean correction only leads to MLEs that are close to the true values. Incorporating the mean and stochastic correction leads also leads a good match between the MLE and the true value, but the profiles widen relative to the profiles for the mean correction only, as previously discussed.

**Figure 11:**
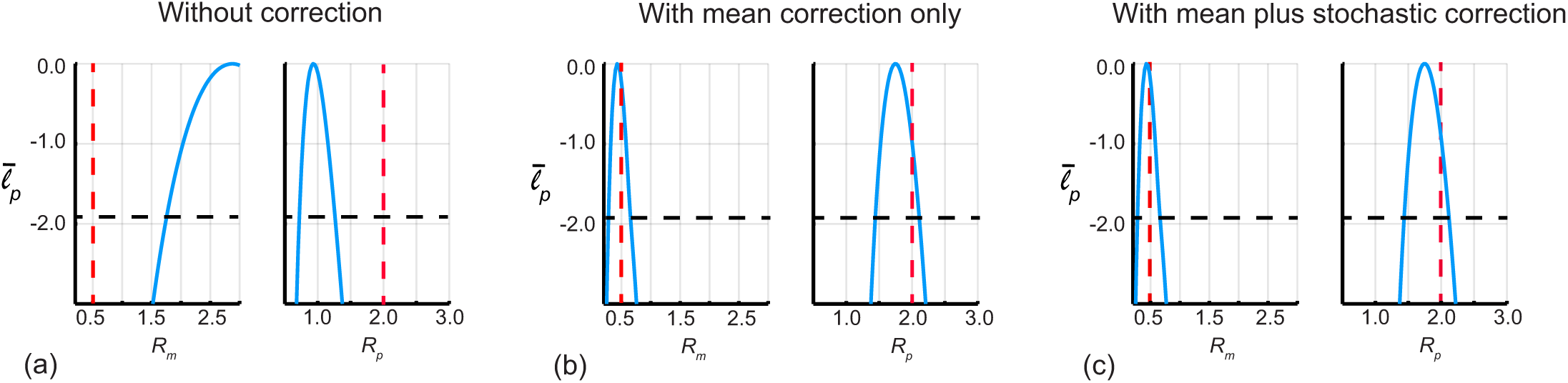
(a)–(c) univariate profile likelihoods, 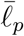, for *R*_*m*_ and *R*_*p*_, as indicated, for the likelihood function without the correction, with the mean correction, and with the mean plus stochastic correction, respectively. The horizontal dashed lines indicates the threshold 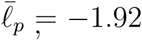 that defines approximate 95% confidence intervals, and the vertical solid red line shows the values used to generate the data. In (a), without the correction, we have 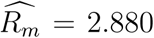, [1.754, 3.000] and 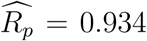, [0.722, 1.266]. In (b), with the mean correction only, we have 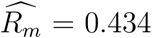, [0.283, 0.654] and 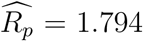, [1.443, 2.110]. In (c), with the mean plus stochastic correction, we have 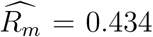, [0.281, 0.661] and 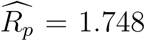, [1.438, 2.120]. All calculations are carried out using *σ*^2^ = 0.05.

## 3 Conclusions and future work

The primary aim of this work was to explore techniques for efficient and reliable parameter inference using stochastic models relevant to cell biology experiments, namely cell proliferation assays and barrier assays. The former experiment is widely adopted to measure cell proliferation rates, and hence the cell doubling time [4, 5, 54], whereas the latter experiment is widely adopted to measure cell migration rates and cell diffusivity *in vitro* [8, 53]. While lattice-based stochastic models are routinely used to simulate such experiments, these simulation models become computationally expensive, particular for inference problems where large numbers of stochastic realisations are required [58]. One way of addressing this computational limitation is to coarse-grain the stochastic models to arrive at an approximate continuum model, often an ODE or PDE model, which is much faster to solve. The limitation of this approach is that the solution of the coarse-grained continuum model does not always accurately predict appropriately averaged data from the discrete model [1, 51]. Here, we illustrate how to combine the advantages of both approaches by using the coarse-grained continuum model together with a model of the discrepancy between the stochastic and continuum models for reliable and efficient parameter inference. We achieve this by using simulation data to sample the discrepancy between the fine stochastic model and the corresponding coarse continuum model and build a statistical model that approximately describes the model error introduced by using the coarse model. Using a series of computational experiments, we demonstrate how to incorporate two levels of correction: the mean correction, or the mean correction together with a stochastic correction. Our numerical experiments show that incorporating the mean correction typically corrects the bias in the likelihood function while broadening the confidence intervals. This broadening of the confidence intervals reflects additional uncertainty introduced through the statistical model of the discrepancy. Incorporating both the mean and stochastic correction leads to a further broadening of the confidence intervals that reflects the additional uncertainty introduced by the approximations.

Within the framework presented here there are other options that could be explored. For example, in both types of computational experiments (i.e. the proliferation assay and the barrier assay), we focus on holding certain properties constant, such as the initial condition, and then treating the parameter estimation and uncertainty quantification problem for two unknown parameters only. This choice was made for pedagogical reasons, but all the analysis and methodology carries across to higher dimensional problems such as modelling a cell proliferation assay where we treat *R*_*p*_, *R*_*d*_ and *C*(0) as unknown quantities to be estimated from the data. Similarly, here we characterise the uncertainty in our estimates by taking a profile likelihood approach and using numerical optimization to construct various profile likelihood functions and then to characterise the uncertainty in our estimates by using threshold values to define appropriate confidence intervals. We chose to characterise uncertainty using profile likelihood-based methods for computational efficiency, but other approaches such as MCMC sampling of the various likelihood functions are a possible alternative [17, 57].

Our methodology for characterising the model error could also have been performed differently. For example, all results in the main document focus on the simplest possible way to sample *θ* by using various uniform distributions. Additional results Section S2 of the Supplementary Material implement the same analysis of the proliferation assay problem except that we sample *θ* using a truncated normal distribution, and analogous results in Section S3 of the Supplementary Material repeat the same computational experiments by sampling *θ* using a Latin hypercube sampling method. Interestingly, regardless of which sampling method we implement, we find that each design leads to virtually indistinguishable results. This is of interest since we anticipate that the choice of sampling method could be important for other applications, especially higher-dimensional problems. Additional results in Section S4 of the Supplementary Material quantitatively examine the assumption that the distribution of *ε* is approximately normal for each sampling method. These results indicate that this assumption is reasonable during the early time where the population is growing, but our data suggests that *ε* is not normally distributed at later time as the population approaches an equilibrium density. Alternative meta-models of the model error, such as Gaussian processes, neural networks, or support vector machines could also be considered [12, 15].

Future work might also extend the approach taken here by working with different stochastic models that incorporate additional biological detail. For example, here we work with the simplest possible approach with a stochastic model where all individuals are treated as identical, but many biological experiments involve co-culture techniques where different cell types are grown together to explore how various cell types interact [16, 61]. Modelling this kind of experiment is straightforward in a stochastic framework but leads to a more computationally demanding simulation algorithm with more parameters to be estimated [61]. Another extension would be to consider modelling how agents interact with each other, for example through modelling cell-to-cell adhesion [9] or different types of cell-to-cell attraction/repulsion mechanisms [48]. This approach is relevant for modelling various types of experiments involving adhesive cancer cells, and incorporating adhesion into the mathematical model is of interest because describing such mechanisms introduces additional parameters to be estimated. Furthermore, coarse-grained PDE models of discrete stochastic models that incorporate adhesion often fail to accurately describe the underlying stochastic model [9]. Therefore, in this instance, the methodology outlined in this work could be used to significantly extend the usefulness of coarse-grained models that might ordinarily be overlooked due to accuracy considerations.

## Supporting information

Additional results and discussion

## Acknowledgements

This work is supported by the Australian Research Council (DP200100177). We thank three anonymous referees for their helpful suggestions.

## Contributions

All authors conceived and designed the study; MJS performed all numerical simulations. All authors drafted the article and gave final approval for publication.

## Data Availability

All algorithms and software required to reproduce the results in this work are available on GitHub.

## Competing Interests

We have no competing interests.

## Notes

https://github.com/ProfMJSimpson/ModelError

### Competing Interest Statement

The authors have declared no competing interest.

## References

[1] Baker RE, Simpson MJ (2010) Correcting mean-field approximations for birth-death-movement processes. Physical Review E. 82, 041905. 10.1103/PhysRevE.82.041905.

[2] Binny RN, James A, Plank MJ (2015) Spatial moment dynamics for collective cell movement incorporating a neighbour-dependent directional bias. Journal of the Royal Society Interface. 12, 20150228. 10.1098/rsif.2015.0228.

[3] Bolker B, Pacala SW (1997) Using moment equations to understand stochastically driven spatial pattern formation in ecological systems. Theoretical Population Biology. 52, 179–197. 10.1006/tpbi.1997.1331.

[4] Bosco DB, Kenworthy R, Zorio DAR, Sang Q-XA (2015) Human mesenchymal stem cells are resistant to paclitaxel by adopting a non-proliferative fibroblastic state. PLOS ONE. 10, e0128511. 10.1371/journal.pone.0128511.

[5] Browning AP, McCue SW, Simpson MJ (2017) A Bayesian computational approach to explore the optimal duration of a cell proliferation assay. Bulletin of Mathematical Biology. 79, 1888–1906. 10.1007/s11538-017-0311-4.

[6] Bruna M, Chapman SJ (2012) Excluded-volume effects in the diffusion of hard spheres. Physical Review E. 85, 011103. 10.1103/PhysRevE.85.011103.

[7] Cox DR, Wermuth, N (1996) Multivariate dependencies: Models, analysis and interpretation. Chapman and Hall/CRC.

[8] Decaestecker C, Debeir O, Van Ham P, Kiss R (2006) Can anti-migratory drugs be screen in vitro? A review of 2D and 3D assays for the quantitative analysis of cell migration. Medicinal Reviews Research. 27, 149–176. 10.1002/med.20078.

[9] Deroulers C, Aubert M, Badoual M, Grammaticos B (2009) Modeling tumor cell migration: from microscopic to macroscopic models. Physical Review E. 79, 031917. 10.1103/PhysRevE.79.031917.

[10] Eaton ML (2007) Multivariate statistics: A vector space approach. Institute of Mathematical Statistics Lecture Notes - Monograph Series, Volume 53.

[11] Fisher RA (1937) The wave of advance of advantageous genes. Annals of Eugenics. 7:355–369.

[12] Forrester A, Sobester A, Keane A (2008) Engineering Design via Surrogate Modelling: a Practical Guide. John Wiley & Sons

[13] Frasca M, Sharkey KJ (2016) Discrete-time moment closure for epidemic spreading in populations of interacting individuals. Journal of Theoretical Biology. 399, 13–21. 10.1016/j.jtbi.2016.03.024.

[14] Gillespie DT (1977) Exact stochastic simulation of coupled chemical reactions. The Journal of Chemical Physics. 81, 2340–2361. 10.1021/j100540a008.

[15] Gramacy, RB (2020) Surrogates: Gaussian process modeling, design, and optimization for the applied sciences. Chapman and Hall/CRC.

[16] Haridas P, Penington CJ, McGovern JA, McElwain DLS, Simpson MJ (2017) Quantifying rates of cell migration and cell prolfieration in co-culture barrier assays reveals how skin and melanoma cells interact during melanoma spreading and invasion. Journal of Theoretical Biology. 423, 13–25. 10.1016/j.jtbi.2017.04.017.

[17] Hines KE, Middendorf TR, Aldrich RW (2014) Determination of parameter identifiability in non-linear biophysical models: A Bayesian approach. Journal of General Physiology. 143, 401–416. 10.1085/jgp.201311116.

[18] Jiang Q, Fu X, Yan S, Li R, Du W, Cao Z, Qian F, Grima R (2021) Neural network aided approximation and parameter inference of non-Markovian models of gene expression. Nature Communications. 12, 2618. 10.1038/s41467-021-22919-1.

[19] Johnson SG (2021) The NLopt module for Julia. Retrieved February 2022 NLopt

[20] Kaipio, J, Somersalo, E (2006) Statistical and computational inverse problems (Vol. 160). Springer.

[21] Kaipio J, Somersalo E (2007) Statistical inverse problems: Discretisation, model reduction and inverse crimes. Journal of Computational and Applied Mathematics. 198, 493–504. 10.1016/j.cam.2005.09.027.

[22] Kaipio J, Kolehmainen, V (2013) Approximate marginalization over modeling errors and uncertainties in inverse problems. Book chapter in Bayesian Theory and Applications. 644–672. 10.1093/acprof:oso/9780199695607.003.0032.

[23] Kennedy MC, O’Hagan A (2000) Predicting the output from a complex computer code when fast approximations are available. Biometrika. 87, 1–13. 10.1093/biomet/87.1.1.

[24] Kennedy MC, O’Hagan A (2001) Bayesian calibration of computer models. Journal of the Royal Statistical Society Series B: Statistical Methdology. 63, 425–464. 10.1111/1467-9868.00294.

[25] Koehler JR, Owen AB (1996). Computer experiments. Handbook of statistics, 13, 261–308.

[26] Kolmogorov AN, Petrovskii PG, Piskunov NS (1937) A study of the diffusion equation with increase in the amount of substance, and its application to a biological problem. Moscow University Mathematics Bulletin. 1, 1–26.

[27] Law R, Murrell DJ, Dieckmann U (2003) Population growth in space and time: Spatial logistic equations. Ecology. 84, 22–22. http://www.jstor.org/stable/3108013.

[28] Lei CL, Ghosh S, Whittaker DG, Aboelkassem Y, Beattie KA, Cantwell CD, Delhaas T, Houston C, Montes Novaes G, Panfilov AV, Pathmanathan P, Riabiz M, Weber dos Santos R, Walmsley J, Worden K, Mirams GR, Wilkinson RD (2020). Considering discrepancy when calibrating a mechanistic electrophysiology model. Philosophical Transactions of the Royal Society A: Mathematical, Physical and Engineering Sciences. 378, 20190349. 10.1098/rsta.2019.0349.

[29] Liang CC, Park A, Guan, JL. In vitro scratch assay: a convenient and inexpensive method for analysis of cell migration in vitro. Nature Protocols. 2, 329–333. 10.1038/nprot.2007.30.

[30] Lipshtat A, Loinger A, Balaban NQ, Biham O (2006) Genetic toggle switch without cooperative binding. Physical Review Letters. 96, 188101. 10.1103/PhysRevLett.96.188101.

[31] Macfarlane FR, Chaplain MAJ, Lorenzi T (2019) A stochastic individual-based model to explore the role of spatial interactions and antigen recognition in the immune response against solid tumours. Journal of Theoretical Biology. 480, 43–55. 10.1016/j.jtbi.2019.07.019.

[32] Macklin P, Edgerton ME, Thompson AM, Cristini V (2012) Patient-calibrated agent-based modelling of ductal carcinoma in situ (DCIS): From microscopic measurements to macroscopic predictions of clinical progression. Journal of Theoretical Biology. 301, 122–140. 10.1016/j.jtbi.2012.02.002.

[33] Maclaren OJ, Nicholson R, Bjarkason EK, O’Sullivan JP, O’Sullivan ML (2020) Incorporating posterior-informed approximation errors into a hierarchical framework to facilitate out-of-the-box MCMC sampling for geothermal inverse problems and uncertainty quantification. Water Resources Research. 56, e24396. 10.1029/2018WR024240.

[34] Macnamara CK, Mitchell EI, Chaplain MAJ (2019) Spatial-stochastic modelling of synthetic gene regulatory networks. Journal of Theoretical Biology. 468, 27–44. 10.1016/j.jtbi.2019.02.003.

[35] Maini PK, McElwain DLS, Leavesley DI (2004) Travelling wave model to interpret a woundhealing cell migration assay for human peritoneal mesothelial cells. Tissue Engineering. 10, 475–482. 10.1089/107632704323061834.

[36] Maini PK, McElwain DLS, Leavesley D (2004) Travelling waves in a wound healing assay. Applied Mathematics Letters. 17, 575–580. 10.1016/S0893-9659(04)90128-0.

[37] McKay, MD (1979) A comparison of three methods for selecting values of input variables in the analysis of output from a computer code. Technometrics. 21(2), 239–245.

[38] Mort RL, Ross RJH, Hainey KJ, Harrison OJ, Keighren MA, Landini G, Baker RE, Painter KJ, Jackson IJ, Yates CA (2016) Reconciling diverse mammalian pigmentation patterns with a fundamental mathematical model. Nature Communications. 7, 1–13. 10.1038/ncomms10288.

[39] Murray JD (2002) Mathematical Biology I: An Introduction. Third edition. Springer, New York.

[40] Myers, RH, Montgomery, DC, Anderson-Cook, CM (2016) Response surface methodology: process and product optimization using designed experiments. John Wiley & Sons.

[41] Owen, AB (1992). A central limit theorem for Latin hypercube sampling. Journal of the Royal Statistical Society: Series B (Methodological). 54(2), 541–551.

[42] Paun LM, Colebank MJ, Olufsen MS, Hill NA, Husmeier D (2020) Assessing model mismatch and model selection in a Bayesian uncertainty quantification analysis of a fluid-dynamics model of pulmonary blood circulation. Journal of the Royal Society Interface. 17, 20200886. 10.1098/rsif.2020.0886

[43] Pawitan Y (2013) In all likelihood: Statistical modelling and inference using likelihood. Oxford University Press, Oxford.

[44] Powell MJD (2009). The BOBYQA algorithm for bound constrained optimization without derivatives. Technical report, Department of Applied Mathematics and Theoretical Physics, Cambridge, England

[45] Raue A, Kreutz C, Maiwald T, Bachmann J, Schilling M, Klingmüller U, Timmer J (2009) Structural and practical identifiability analysis of partially observed dynamical models by exploiting the profile likelihood. Bioinformatics. 25, 123–12. 10.1093/bioinformatics/btp358.

[46] Royston P (2007) Profile likelihood for estimation and confidence intervals. The Stata Journal. 7, 376–387. 10.1177/1536867X0700700305.

[47] Santner, TJ, Williams, BJ, Notz, WI, Williams, BJ (2003) The design and analysis of computer experiments. New York: Springer.

[48] Simpson MJ, Landman KA, Hughed BD, Fernando AE (2010) A model for mesoscale patterns in motile populations. Physica A: Statistical Mechanics and its Applications. 389, 1412–1424. 10.1016/j.physa.2009.12.010.

[49] Simpson MJ, Landman KA, Hughed BD (2010) Cell inavasion with proliferation mechanisms motivated by time-lapse data. Physica A: Statistical Mechanics and its Applications. 389, 3779–3790. 10.1016/j.physa.2010.05.020.

[50] Simpson MJ, Binder BJ, Haridas P, Wood BK, Treloar KK, McElwain DLS, Baker RE (2013) Experimental and modelling investigation of monolayer development with clustering. Bulletin of Mathematical Biology. 75, 871–889. 10.1007/s11538-013-9839-0.

[51] Simpson MJ, Sharp JA, Baker RE (2014) Distinguishing between mean-field, moment dynamics and stochastic description of birth-death-movement processes. Physica A: Statistical Mechanics and its Applications. 395, 236–246. 10.1016/j.physa.2013.10.026.

[52] Simpson MJ, Browning AP, Warne DJ, Maclaren OJ, Baker RE (2022) Parameter identifiability and model selection for sigmoid population growth models. Journal of Theoretical Biology. 535, 110998. 10.1016/j.jtbi.2021.110998.

[53] Treloar KK, Simpson MJ, McElwain DLS, Baker RE (2014) Are in vitro estimates of cell diffusivity and cell proliferation rate sensitive to assay geometry? Journal of Theoretical Biology. 356, 71–84. 10.1016/j.jtbi.2014.04.026.

[54] Tremel A, Cai A, Tirtaatmadja N, Hughes DB, Stevens GW, Landman KA, O’Connor AJ (2009) Cell migration and proliferation during monolayer formation and wound healing. Chemical Engineering Science. 64, 247–253. 10.1016/j.ces.2008.10.008.

[55] Vellela M, Qian H (2007) A Quasistationary analysis of a stochastic chemical reaction: Keizer’s paradox. Bulletin of Mathematical Biology. 68, 1727–1746. 10.1007/s11538-006-9188-3.

[56] Vittadello ST, McCue SW, Gunasingh G, Haass NK, Simpson MJ (2018) Mathematical models for cell migration with real-time cell cycle dynamics. Biophysical Journal. 114, 1241–1253. 10.1016/j.bpj.2017.12.041.

[57] Villaverde AF, Pathirana D, Fröhlich F, Hasenauer J, Banga JR (2022) A protocol for dynamic model calibration. Briefings in Bioinformatics. 23, 1–19. 10.1093/bib/bbab387.

[58] Vo BN, Drovandi CC, Pettitt AN, Simpson MJ (2005) Quantifying uncertainty in parameter estimates for stochastic models of collective cell spreading using approximate Bayesian computation. Mathematical Biosciences. 263, 133–142. 10.1016/j.mbs.2015.02.010.

[59] Wooldridge, J. M. (2010). Econometric analysis of cross section and panel data. MIT press.

[60] Xu T, Valocchi AJ, Ye M, Liang F (2017) Quantifying model structural error: Efficient Bayesian calibration of a regional groundwater flow model using surrogates and a data-driven error model. Water Resources Research. 3, 4084–4105. 10.1002/2016WR019831.

[61] Zhang D, Brinas IM, Binder BJ, Landman KA, Newgreen DF (2010) Neural crest regionalisation for enteric nervous system formation: Implications for Hirschsprung’s disease and stem cell therapy. Developmental Biology. 339, 280–294. 10.1016/j.ydbio.2009.12.014.

[62] Ziaeipoor H, Taylor M, Pandy M, Martelli S (2019) A novel training-free method for real-time prediction of femoral strain. Journal of Biomechanics. 86, 110–116. 10.1016/j.jbiomech.2019.01.057.

