## Additional results and discussion for "Reliable and efficient parameter estimation using approximate continuum limit descriptions of stochastic models"

### Contents

**S1 Numerical methods for the partial differential equation model** **2**

**S2 Sampling $\theta$ using a truncated multivariate normal distribution** **3**

**S3 Sampling $\theta$ using a Latin hypercube design** **5**

**S4 Working assumption that $\varepsilon$ is normally distributed** **8**

### S1 Numerical methods for the partial differential equation model

We solve Equation (5) by first discretising the spatial terms on a uniform mesh, with mesh spacing $h > 0$, so that $C(x_k, t) = C(k)$ for $k = 1, 2, 3, \ldots, K$, where $x_k = (k-1)h$. This gives,

$$\begin{aligned} 0 &= C(1) - C(2), \\ \frac{\mathrm{d}C(k)}{\mathrm{d}t} &= \frac{D}{h^2}\left[C(k-1) - 2C(k) + C(k+1)\right] + r_1 C(k)\left[1 - C(k)\right] - r_2 C(k), \quad k = 2, 3, 4, \ldots, K-1 \\ 0 &= C(K) - C(K-1). \end{aligned} \tag{S1}$$

We then solve this system of differential algebraic equations using an explicit method with automatic truncation error control in Julia [1, 2]. All results reported are obtained with $h = 1.0$, and we obtain visually indistinct results with $h = 0.5$, suggesting that our results are grid-independent.

### S2 Sampling $\theta$ using a truncated multivariate normal distribution

Here we briefly return to the question of simulation and inference of a cell proliferation assay reported previously in Figures 4–7 where we sampled $\theta$ using a uniform distribution. We now repeat the simulations in Figure 4 by sampling $\theta$ from a truncated normal distribution (constrained to be non-negative), as illustrated in Figure S1(a). Results in Figure S1(b)–(d) confirm that sampling $\theta$ in this way allows us to perform accurate simulations with the continuum limit model for the same parameter combinations as we explored in Figure 4.

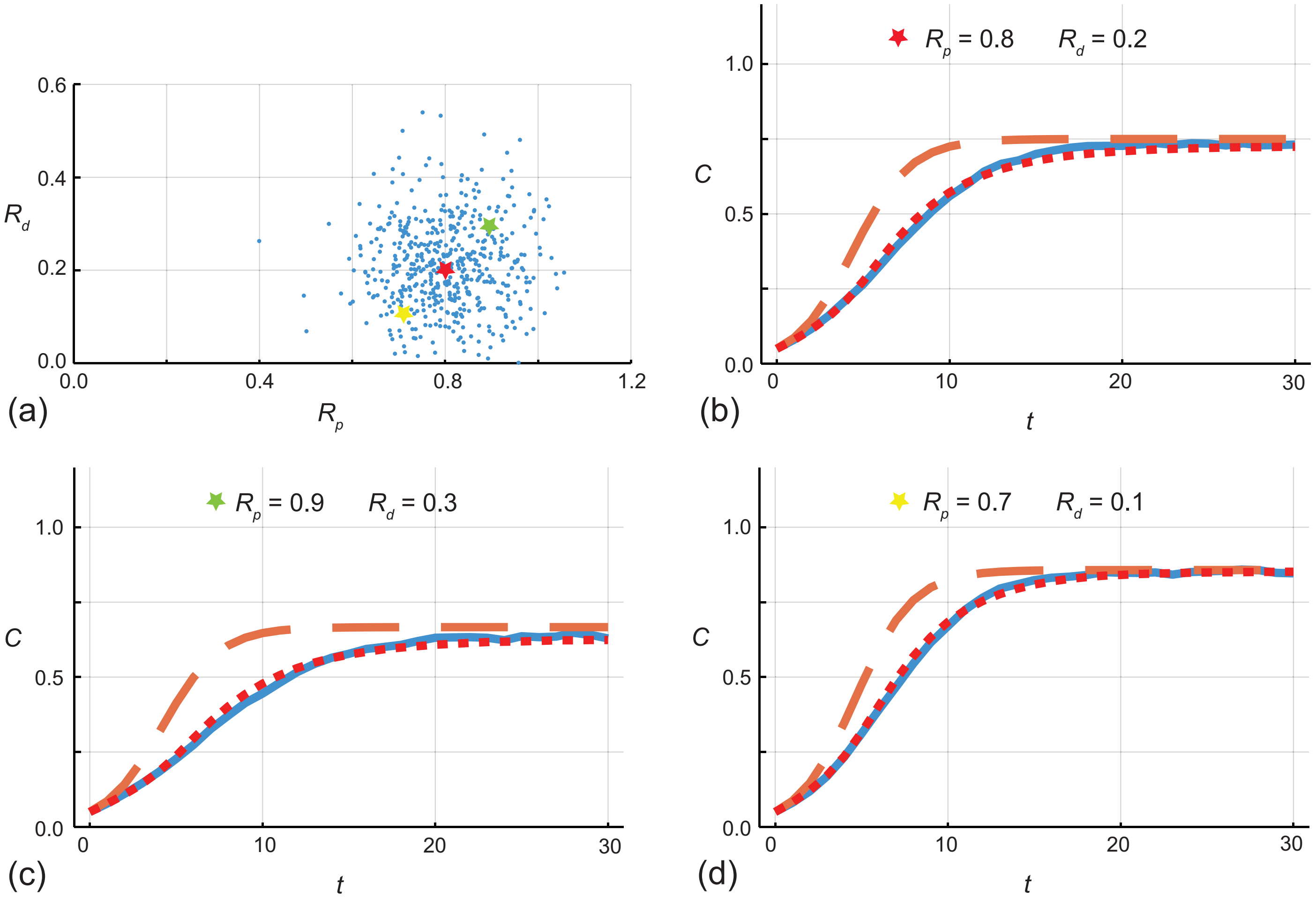


Figure S1: (a) $\mathcal{N}_d = 500$ samples of pairs of $[R_p, R_d]$ drawn from a truncated bivariate normal distribution with mean $\mu = [0.8, 0.2]^\top$ and diagonal covariance $\Sigma = \text{diag}(0.1, 0.1)$ used to construct the error model. The bivariate normal distribution is truncated so that $R_p > 0$ and $R_d > 0$. (b)–(d) Comparison of $C(t \mid \theta)$ for the fine (stochastic) model (solid blue), the coarse (continuum) model (dashed orange) and the coarse model plus mean correction term (dotted red) for $[R_p, R_d] = [0.8, 0.2]$, $[0.9, 0.3]$ and $[0.7, 0.1]$, respectively. Stochastic simulations are performed on a lattice with $I = J = 100$, with $C(0 \mid \theta) = 0.1$ and $R_m = 1$. Stochastic simulation data is recorded at $t = 1, 2, \ldots, 29, 30$, giving $\mathcal{N}_t = 30$.

With estimates of the correction term obtained using the sampling in Figure S1 we compute the likelihood, both without and with the correction, as given in Figure S2. Just like in Figure 6 we see that without the correction term the 95% confidence region does not contain the true value, but when we incorporate the mean correction only or the mean plus stochastic correction terms the true value is centered within the 95% confidence region. Profile likelihoods, both without and with the correction terms, are given in Figure S3.

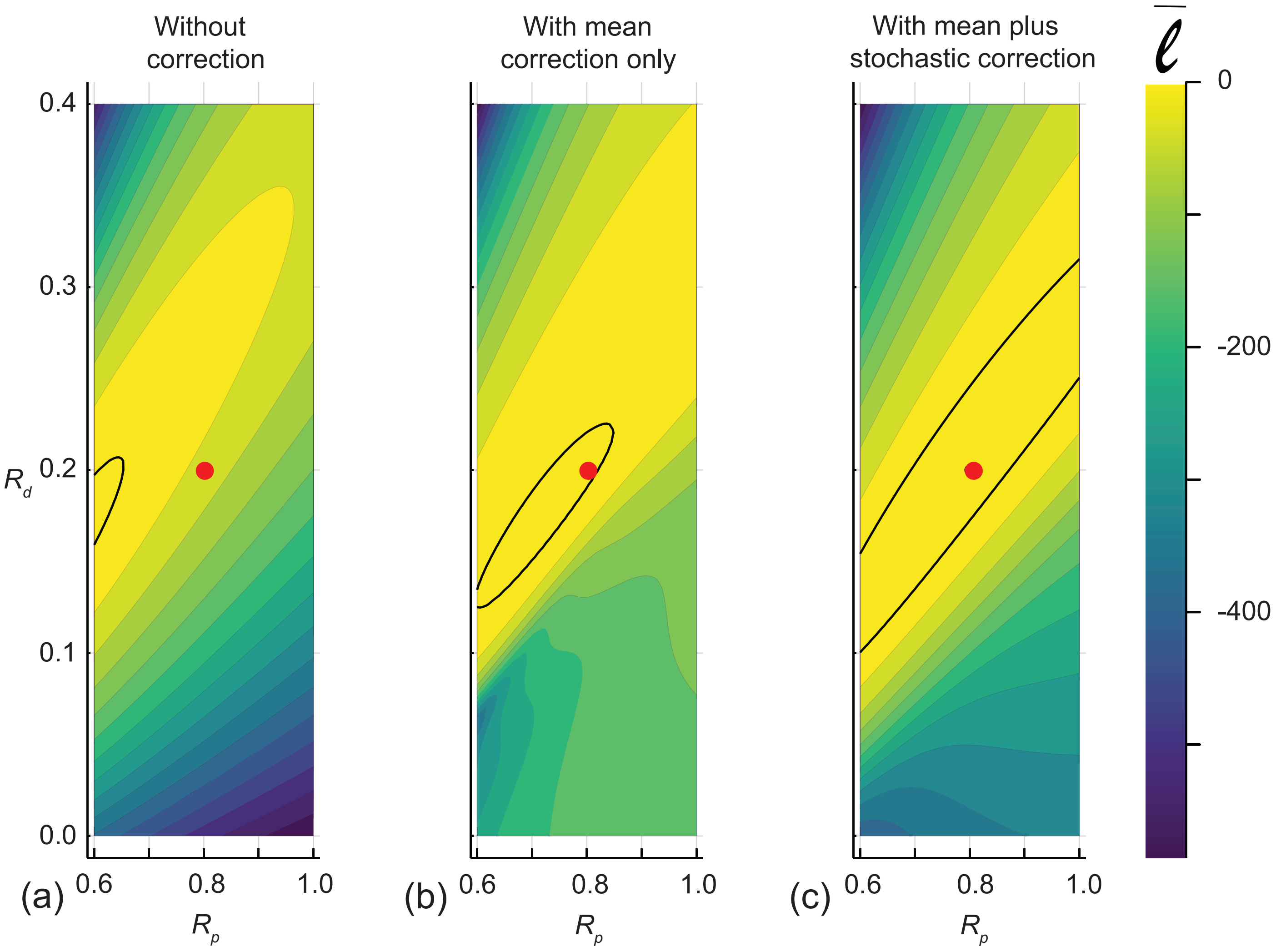


Figure S2: (a)–(c) show the normalised log likelihood as a function of $[R_p, R_d]$ for the coarse approximate model without any correction, with mean correction only, and with mean plus stochastic correction, respectively. Each heat map is constructed by evaluating the likelihood across a $40 \times 40$ uniform discretisation of the parameter space. The outline of the region where $\bar{\ell} = -3.00$ defines the approximate 95% confidence region and the location of the values used to generate the data, $[R_p, R_d] = [0.8, 0.2]$, is superimposed on each plot. All calculations are carried out using $\sigma^2 = 0.05$.

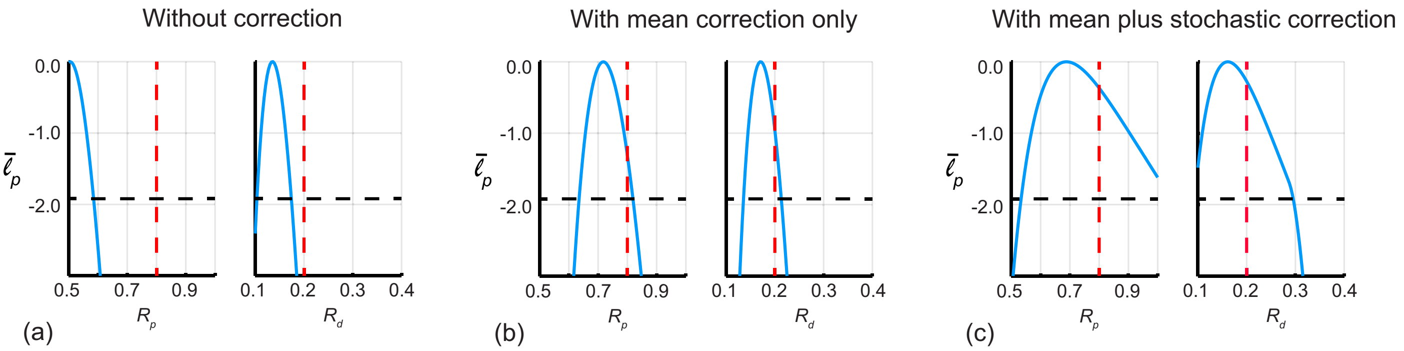


Figure S3: (a)–(c) univariate profile likelihoods, $\bar{\ell}_p$, for $R_m$ and $R_p$, as indicated, for the likelihood function without the correction, with the mean correction, and with the mean plus stochastic correction, respectively. The horizontal dashed lines indicates the threshold $\bar{\ell}_p = -1.92$ that defines approximate 95% confidence intervals, and the vertical solid red line shows the value used to generate the data. In (a), without the correction, we have $\widehat{R_p} = 0.505$, $[0.500, 0.585]$ and $\widehat{R_d} = 0.135$, $[0.103, 0.174]$. In (b), with the mean correction only, we have $\widehat{R_p} = 0.712$, $[0.640, 0.819]$ and $\widehat{R_d} = 0.170$, $[0.140, 0.212]$. In (c), with the mean plus stochastic correction, we have $\widehat{R_p} = 0.688$, $[0.533, 1.000]$ and $\widehat{R_d} = 0.161$, $[0.100, 0.295]$. All calculations are carried out using $\sigma^2 = 0.05$.

### S3 Sampling $\theta$ using a Latin hypercube design

Here we briefly return to the question of simulation and inference of a cell proliferation assay reported previously in Figures 4–7 where we sampled $\theta$ using a uniform distribution. We now repeat the simulations in Figure 4 by sampling $\theta$ using a Latin hypercube design [3], as illustrated in Figure S4(a). Results in Figure S4(b)–(d) confirm that sampling $\theta$ in this way allows us to perform accurate simulations with the continuum limit model for the same parameter combinations explored in Figure 4.

With estimates of the correction term obtained using the sampling in Figure S4, we now compute the likelihood functions reported in Figure S5 and the various univariate profile likelihood functions in Figure S6.

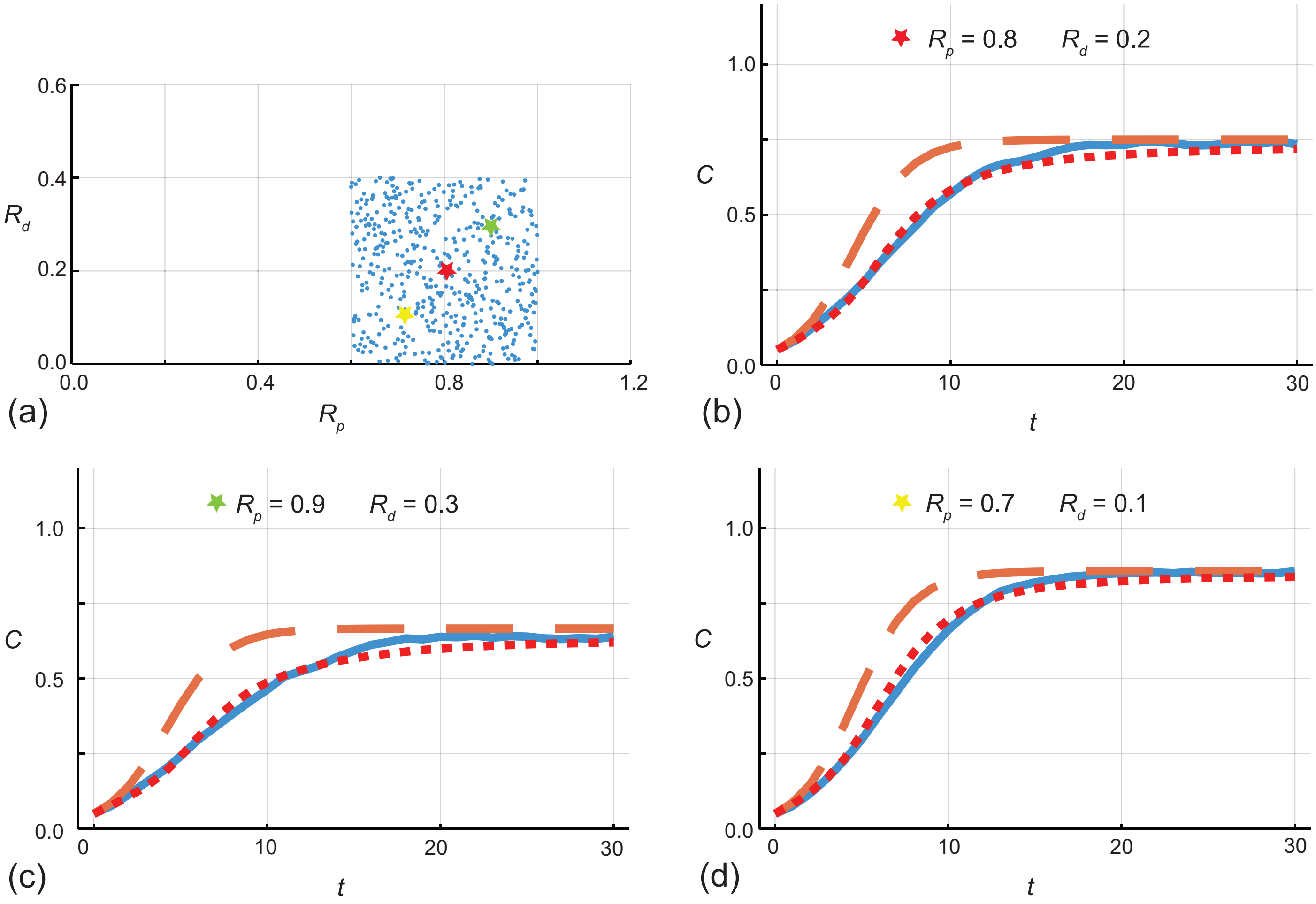


Figure S4: (a) $\mathcal{N}_d = 500$ samples of pairs of $[R_p, R_d]$ drawn using Latin hypercube sampling in the region $R_p \in [0.7, 0.9]$ and $R_d \in [0.0, 0.4]$. (b)–(d) Comparison of $C(t \,|\, \theta)$ for the fine (stochastic) model (solid blue), the coarse (continuum) model (dashed orange) and the coarse model plus mean correction term (dotted red) for $[R_p, R_d] = [0.8, 0.2]$, $[0.9, 0.3]$ and $[0.7, 0.1]$, respectively. Stochastic simulations are performed on a lattice with $I = J = 100$, with $C(0 \,|\, \theta) = 0.1$ and $R_m = 1$. Stochastic simulation data is recorded at $t = 1, 2, \ldots, 29, 30$, giving $\mathcal{N}_t = 30$.

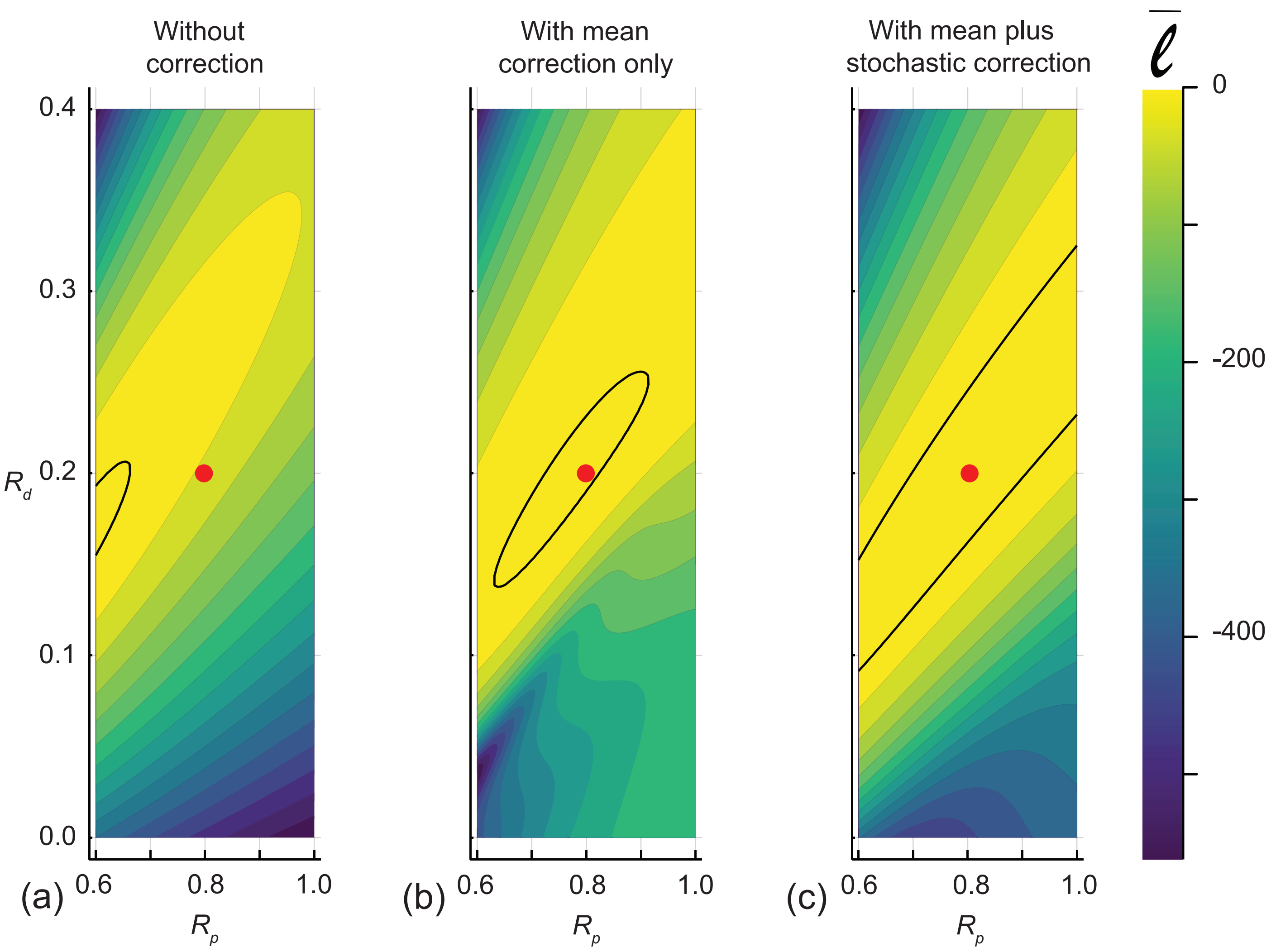


Figure S5: (a)–(c) show the normalised log likelihood as a function of $[R_p, R_d]$ for the coarse approximate model without any correction, with mean correction only, and with mean plus stochastic correction, respectively. Each heat map is constructed by evaluating the likelihood across a $40 \times 40$ uniform discretisation of the parameter space. The outline of the region where $\bar{\ell} = -3.00$ defines the approximate 95% confidence region and the location of the values used to generate the data, $[R_p, R_d] = [0.8, 0.2]$, is superimposed on each plot. All calculations are carried out using $\sigma^2 = 0.05$.

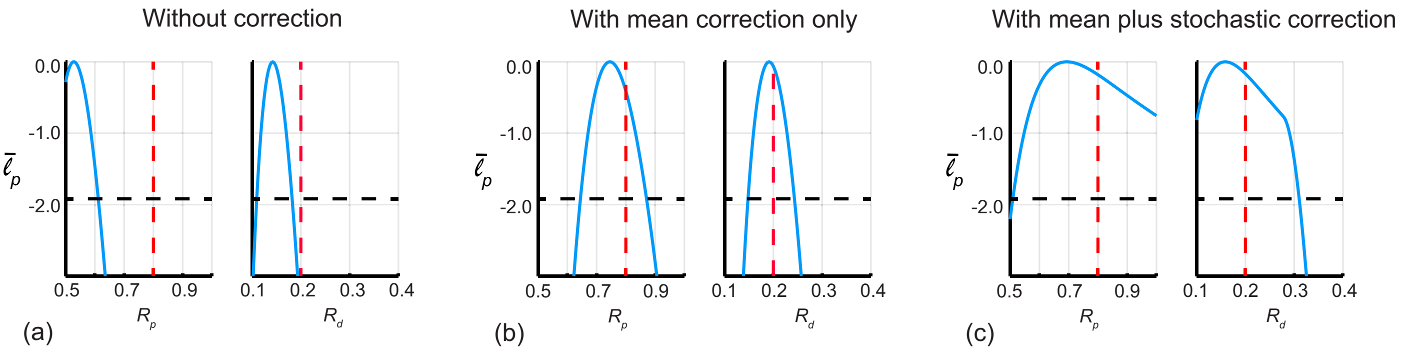


Figure S6: (a)–(c) univariate profile likelihoods, $\bar{\ell}_p$, for $R_m$ and $R_p$, as indicated, for the likelihood function without the correction, with the mean correction, and with the mean plus stochastic correction, respectively. The horizontal dashed lines indicates the threshold $\bar{\ell}_p = -1.92$ that defines approximate 95% confidence intervals, and the vertical solid red line shows the values used to generate the data. In (a), without the correction, we have $\widehat{R_p} = 0.528$, $[0.500, 0.612]$ and $\widehat{R_d} = 0.142$, $[0.109, 0.182]$. In (b), with the mean correction only, we have $\widehat{R_p} = 0.745$, $[0.644, 0.871]$ and $\widehat{R_d} = 0.192$, $[0.148, 0.243]$. In (c), with the mean plus stochastic correction, we have $\widehat{R_p} = 0.694$, $[0.508, 1.000]$ and $\widehat{R_d} = 0.159$, $[0.100, 0.310]$. All calculations are carried out using $\sigma^2 = 0.05$.

### S4 Working assumption that $\varepsilon$ is normally distributed

An implicit assumption in our multivariate linear regression procedure is that $\varepsilon$ is approximately normally distributed, and we now examine the validity of this assumption. Data in Figures S7–S9 explore data at $t = 5, 10$ and 15 for simulations of the proliferation assay when $\theta$ is sampled using the three different designs considered in this work. We choose to examine the distributions of $\varepsilon$ at these particular times since they are three equally-spaced time points during the population growth process before the density reaches equilibrium at late time, as indicated by the evolution of $C(t)$ in Figure 4. Histograms and Q-Q plots provide a visual assessment of the assumption that $\varepsilon$ is normally distributed and for each design we see that the assumption becomes less valid as time increases. Regardless of this trend, our results confirm that the statistical meta-model of the model discrepancy performs well, even at late time when $\varepsilon$ is not normally distributed

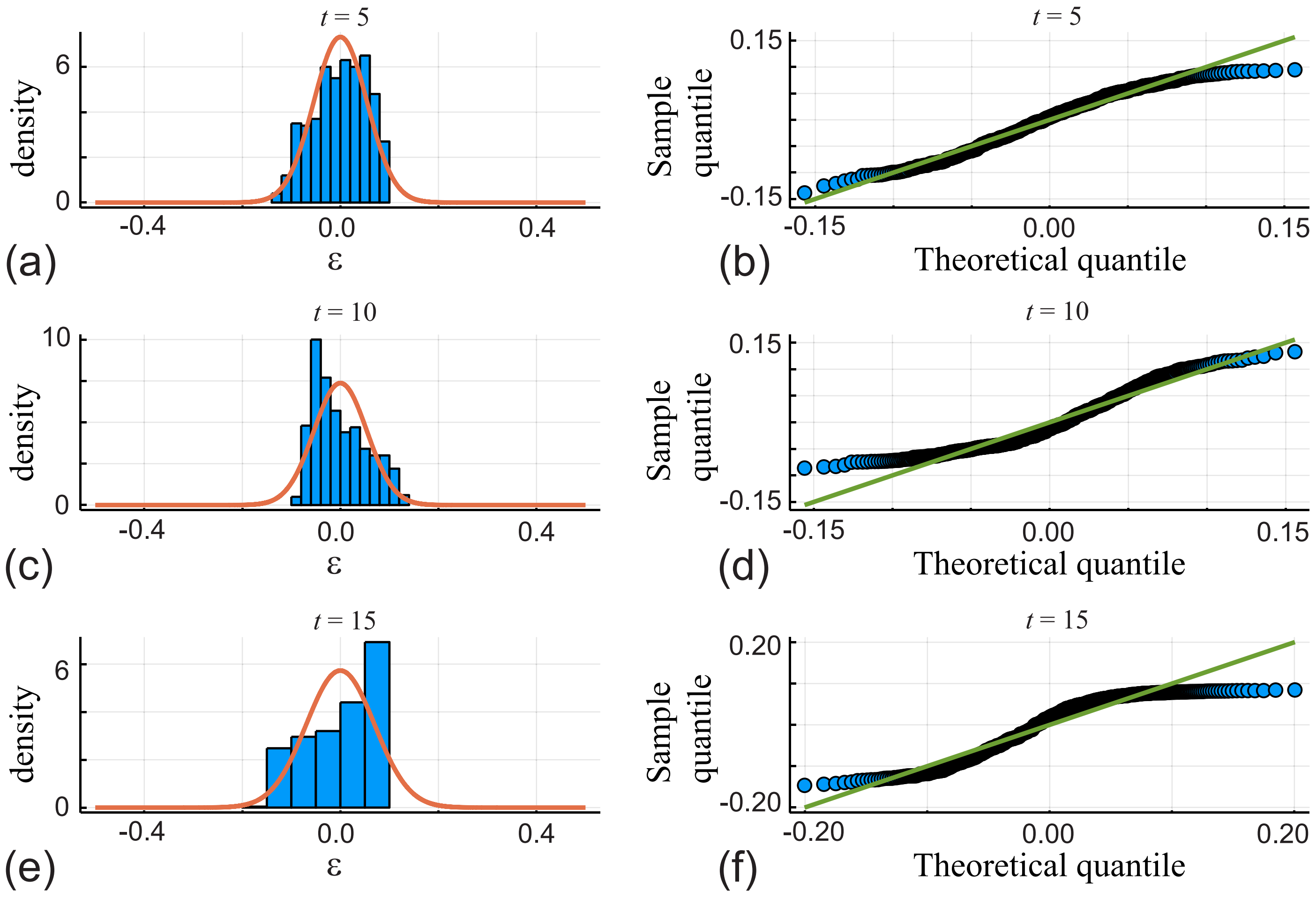


Figure S7: (a)–(b), (c)–(d), (e)–(f) histograms of $\varepsilon$ data and associated Q-Q plots for $t = 5, 10$ and 15 when $\theta$ is sampled from the uniform distribution, as in Figure 4, using $\mathcal{N}_d = 500$ samples. In (a), (c) and (e) $\varepsilon$ data is shown as histograms superimposed with a univariate Gaussian probability density function, with mean and variance estimated from that data. In (b), (d) and (f) we show Q-Q plots comparing the sample quantile from the data with the theoretical quantile assuming the data is normally distributed.

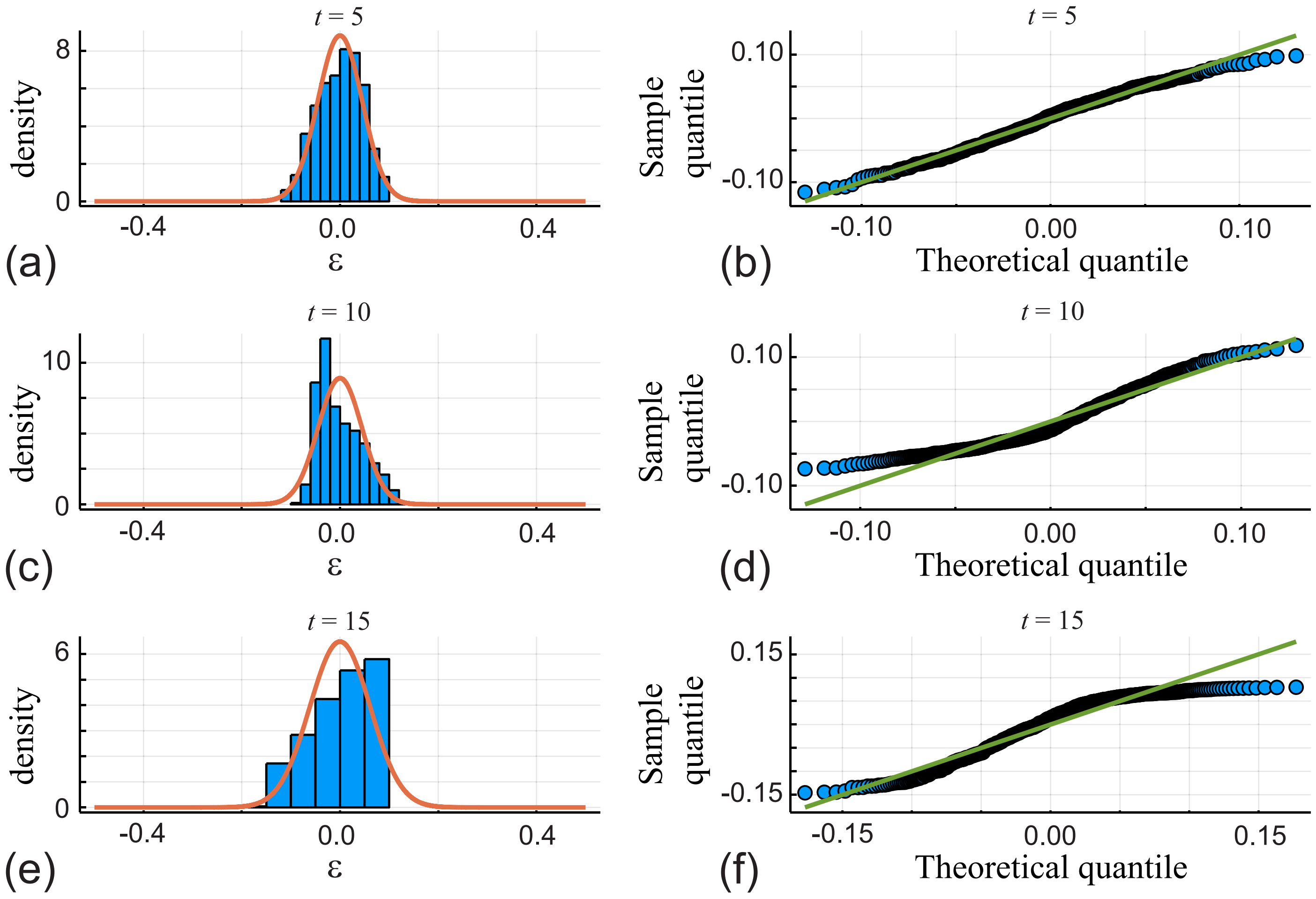


Figure S8: (a)–(b), (c)–(d), (e)–(f) histograms of $\varepsilon$ data and associated Q-Q plots for $t = 5, 10$ and 15 when $\theta$ is sampled from the truncated bivariate normal distribution, as in Figure S1, using $\mathcal{N}_d = 500$ samples. In (a), (c) and (e) $\varepsilon$ data is shown as histograms superimposed with a univariate Gaussian probability density function, with mean and variance estimated from that data. In (b), (d) and (f) we show Q-Q plots comparing the sample quantile from the data with the theoretical quantile assuming the data is normally distributed.

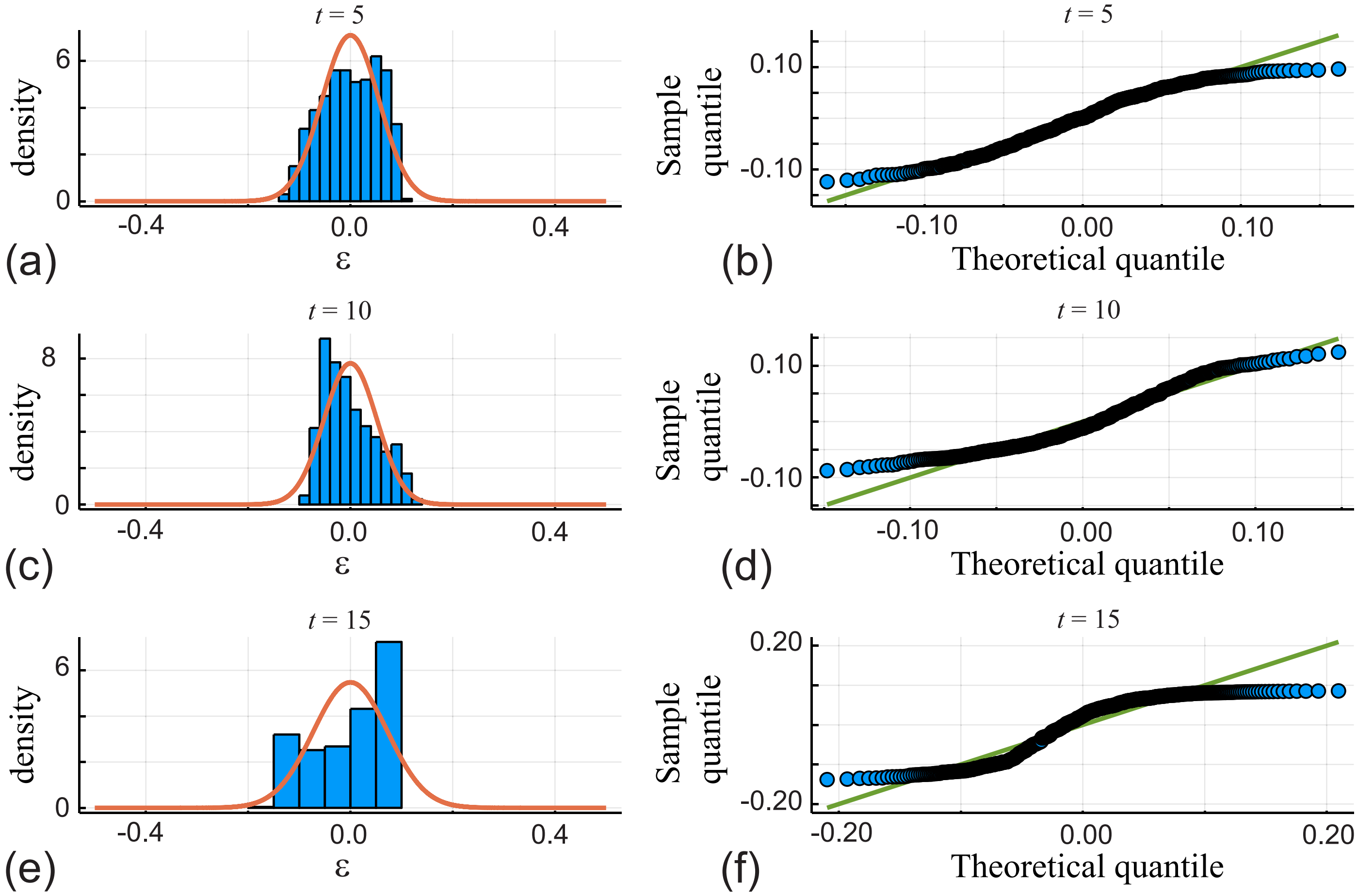


Figure S9: (a)–(b), (c)–(d), (e)–(f) histograms of $\varepsilon$ data and associated Q-Q plots for $t = 5, 10$ and 15 when $\theta$ is sampled from the Latin hypercube design, as in Figure S4, using $\mathcal{N}_d = 500$ samples. In (a), (c) and (e) $\varepsilon$ data is shown as histograms superimposed with a univariate Gaussian probability density function, with mean and variance estimated from that data. In (b), (d) and (f) we show Q-Q plots comparing the sample quantile from the data with the theoretical quantile assuming the data is normally distributed.

### References


[1] Rackauckas C, Nie Q (2017) DifferentialEquations.jl – A performant and feature-rich ecosystem for solving differential equations in julia. Journal of Open Research Software. 5, 15. 10.5334/jors.151.

[2] Tsitouras Ch (2011) Runge–Kutta pairs of order 5(4) satisfying only the first column simplifying assumption. Computers & Mathematics with Applications. 62, 770–775. 10.1016/j.camwa.2011.06.002.

[3] Urquhart M, Ljungskog E, Sebben S (2020) Surrogate-based optimisation using adaptively scaled radial basis functions. Applied Soft Computing. 88, 1568–4946. 10.1016/j.asoc.2019.106050.
